# Fine-tuning of conditional Transformers for the generation of functionally characterized enzymes

**DOI:** 10.1101/2024.08.10.607430

**Authors:** Marco Nicolini, Emanuele Saitto, Ruben Emilio Jimenez Franco, Emanuele Cavalleri, Marco Mesiti, Aldo Javier Galeano Alfonso, Dario Malchiodi, Alberto Paccanaro, Peter N. Robinson, Elena Casiraghi, Giorgio Valentini

## Abstract

We introduce *Finenzyme*, a Protein Language Model (PLM) that employs a multifaceted learning strategy based on transfer learning from a decoder-based Transformer, conditional learning using specific functional keywords, and fine-tuning to model specific Enzyme Commission (EC) categories. Using *Finenzyme*, we investigate the conditions under which fine-tuning enhances the prediction and generation of EC categories, showing a two-fold perplexity improvement in EC-specific categories compared to a generalist model. Our extensive experimentation shows that *Finenzyme* generated sequences can be very different from natural ones while retaining similar tertiary structures, functions and chemical kinetics of their natural counterparts. Importantly, the embedded representations of the generated enzymes closely resemble those of natural ones, thus making them suitable for downstream tasks. Finally, we illustrate how *Finenzyme* can be used in practice to generate enzymes characterized by specific functions using in-silico directed evolution, a computationally inexpensive PLM fine-tuning procedure significantly enhancing and assisting targeted enzyme engineering tasks.

## 1 Introduction

Large Language Models (LLM) trained on biomedical texts, electronic health records, protein, and DNA sequences have achieved unprecedented success in a large range of biomedical domains (Wang et al., 2024a). Protein Language Models (PLM), which leverage the similarity between natural language and “protein language” (Ofer et al., 2021), can learn complex molecular distributions of sequences of amino acids (Flam-Shepherd et al., 2022) in a manner similar to how LLMs model the distribution of word sequences in the context of Natural Language Processing (NLP) (Ferruz and Höcker, 2022). Analogously to LLM models for NLP, PLMs share a similar Transformer architecture (Vaswani et al., 2017) and can learn interactions between amino acid residues, motifs, and domains through the self-attention mechanism, by stacking multiple layer modules (Valentini et al., 2023). Moreover, these models can be directly trained via self-supervised learning from large public domain protein repositories such as UniParc or UniProt (The UniProt Consortium, 2023), without requiring labeled examples.

Encoder-based Transformers have been mainly applied to obtain embedded representations of proteins for a wide spectrum of downstream tasks, ranging from the prediction of long-range contacts and mutational effects (Rives et al., 2021) to the selection of antibodies with improved binding affinity (Hie et al., 2023), as well as secondary structure, remote homology and signal peptide prediction (Brandes et al., 2022). A Multiple Sequence Alignment (MSA) Encoder combined with a bidirectional long short-term memory (BiLSTM) network achieved breakthrough results in the classification of signal peptides and prediction of cleavage site positions (Shen et al., 2024). Decoder-based Transformers are generative models, in the sense that they are able to generate the next amino acid in a sequence, on the basis of the previous ones by using masked self-attention layers. For example, ProtGPT2 can not only generate sequences similar to those of natural proteins but can also generate novel sequences not selected by natural evolution processes (Ferruz and Schmidt, 2022). Although decoder-based Transformers are basically generative models, their last hidden layers provide embedded representations of proteins that can be used for downstream tasks, such as the prediction of mutational effects of functional loss (Ferruz and Höcker, 2022), or scoring protein sequences (Ruffolo and Madani, 2024).

The modeling and generation of functionally characterized proteins is one of the main objectives of protein engineering (Li et al., 2020); by prepending a keyword that functionally characterizes the protein to be generated, conditional Transformers open the way for protein design tailored to specific protein features (Madani et al., 2023). For example, we can condition protein generation on a functional tag for a specific enzymatic reaction and at the same time on another tag for a specific binding domain, generating proteins that can drive a specific biochemical reaction in a specific micro-environment.

Fine-tuning is another technique largely used to improve the performance of pre-trained autoregressive LLMs. According to this approach, pre-trained models undergo a second training step using additional data for a specific task to fine-tune model parameters, sometimes also adding new neural layers on top of the models to specialize them for specific learning tasks (Chung et al., 2024). This is distinct from zero-shot scenarios, where the model generates sequences without any additional training, relying solely on its pre-existing knowledge.

Several studies have demonstrated the effectiveness of fine-tuning in the context of PLMs. For instance, parameter-efficient fine-tuning of PLMs improved protein-protein interactions (Sledzieski et al., 2023); analogously, fine-tuning of pre-trained antibody language models boosted the binding antigen specificity predictions to SARS-CoV2 spike protein and influenza hemagglutinin (Wang et al., 2024b). Lafita et al. (2024) focused on fine-tuning ESM family models, showing that fine-tuning PLMs with deep mutational scanning improves variant effect prediction. By implanting low-rank adaption (LoRA) into ESM2 models, Zeng et al. (2023) were able to improve signal peptide prediction. Fine-tuning the ProtBert-BFD and Prot5-XL-Uniref50 Transformers (Elnaggar et al., 2022) led to a significant improvement in Gene Ontology and Enzyme Commission (EC) number predictions (Wenzel et al., 2024), close to the top-model results of the CAFA3 challenge (Zhou, N., Jiang, Y., Bergquist, T.R. and others, 2019). Exploiting Prot-Bert and ESM-2 models for fine-tuning with new data and adding supervised classifiers on top of the LLMs, Ghazikhani and Butler (2024) improved the classification of ion channels and ion transporters. Our results on the fine-tuning of the ProGen model on the lysozyme family of proteins achieved a statistically significant improvement in terms of accuracy and perplexity (Nicolini et al., 2024), confirming previous results on a similar task (Madani et al., 2023).

Other studies have outlined the drawbacks and limitations of fine-tuning, demonstrating that this approach sometimes fails to achieve the desired results. For instance in the context of biomedical NLP, Tinn et al. (2023) showed that fine-tuning stability and performance depends on the characteristics of the pre-trained model, whose performance cannot always be improved using this technique. In the context of PLMs, Schmirler et al. (2024) demonstrated that finetuning ESM2 and ProstT5 (Heinzinger et al., 2024) models boosts predictions in various tasks, ranging from disorder and mutation effects prediction to sub-cellular location prediction, while no statistically significant improvement can be achieved in other tasks, such as secondary structure prediction.

In this work, we investigate the conditions under which fine-tuning improves the performance of pretrained PLMs applied to enzyme prediction and generation. Enzymes, accelerating chemical reactions by orders of magnitude, play a key role in cellular biology and span fundamental medical and industrial applications, ranging from pharmacology to the reduction of environmental pollution (Walsh and Walsh, 2022; Yeh et al., 2023). We propose a dual learning approach: a) PLM conditional learning and b) fine-tuning of general models pre-trained on a large corpus of proteins, not limited to enzymes only. From this standpoint, our approach follows a different learning strategy than the recently proposed specialized pre-training of conditional Transformers on natural enzymes downloaded from the BRENDA database (Munsamy et al., 2024). Indeed, our proposed *Finenzyme* models combine conditional Transformers pre-trained on large corpora of proteins with well-focused fine-tuning on specific enzymatic categories to generate enzymes that on the one hand maintain the basic properties of natural enzymes and on the other hand explore avenues not probed by natural evolution.

Our primary contributions are summarized as follows:

- We propose a new model, *Finenzyme*, based on a multi-faceted strategy to learn EC (Enzyme Commission) categories of enzymes: PLM transfer learning, conditional learning and finetuning.
- We demonstrate that fine-tuning boosts prediction and generation of specific EC categories, whereas for more general EC categories the improvement is negligible.
- We provide fine-tuned models that generate from scratch enzymes close to natural ones, using only keywords associated with each EC category.
- We show how to generate enzymes “evolved” from natural counterparts through *Finenzyme* to produce new sequences with well-characterized functional properties.
- We characterize the enzymes generated by *Finenzyme* models by comparing the 3D representations of natural and generated enzymes using ESMfold (Lin et al., 2023) and Foldseek (Van Kempen et al., 2024), showing that the primary and tertiary structure of the enzymes generated by *Finenzyme* models resemble those of natural ones.
- We show that the generated enzymes maintain the same functions of their natural counterparts and preserve the chemical kinetics of natural enzymes, sometimes even improving them.
- We characterize the *Finenzyme* generated enzymes by clustering the 3D structure representations of natural and generated enzymes, revealing that the resulting clusters are largely superposed.
- We provide further evidence of the effectiveness of *Finenzyme*, showing that embedded representations of generated and natural enzymes are very similar, independently if obtained from *Finenzyme* or ESM2 models.
- We provide source code to reproduce all experiments, scripts, and tutorials to allow users to fine-tune *Finenzyme* on any EC category or groups of functionally related EC categories.

## 2 Results

In our experiments, we fine-tuned ProGen (Madani et al., 2023), a protein language model with 1.2 billion parameters that was trained on a dataset of 280 million protein sequences from Uniparc, UniprotKB, and SwissProt. ProGen generates an amino acid sequence in an autoregressive way, while prepending functional tags to guide the generation. Fig. S1 (Supplementary Information) provides a high level scheme of the conditional Transformer we used in the experiments.

We fine-tuned ProGen to learn the enzyme characteristics of specific EC classes through two combined learning strategies: a) conditional decoder learning, that is generating enzymes conditioned on a specific EC tag; b) transfer learning and fine-tuning of a pretrained model on specific EC classes (see Material and Methods for more details).

We first show that *Finenzyme* significantly improves ProGen sequence prediction of the most specific EC categories. Then we highlight that the primary and tertiary structures of the enzymes generated by *Finenzyme* are close to natural ones, using ESMFold (Lin et al., 2023), FoldSeek (Van Kempen et al., 2024) and CLEAN (Yu et al., 2023b) tools, and their functions are preserved

We report results obtained by fine-tuning *Finenzyme* on a group of functionally related EC categories belonging to EC 3.8.1, demonstrating that the embedded representations of generated and natural enzymes are similar, independently if generated from *Finenzyme* or ESMFold We also show that clusters of embedded representations of natural and generated enzymes largely superpose, and the same embeddings can be used to accurately classify EC categories.

Finally, we propose how to generate functionally characterized enzymes through *Finenzyme* directed evolution from natural sequences. Fig. 1 provides a schematic summary of the main computational tasks performed in this work.

**Figure 1:**
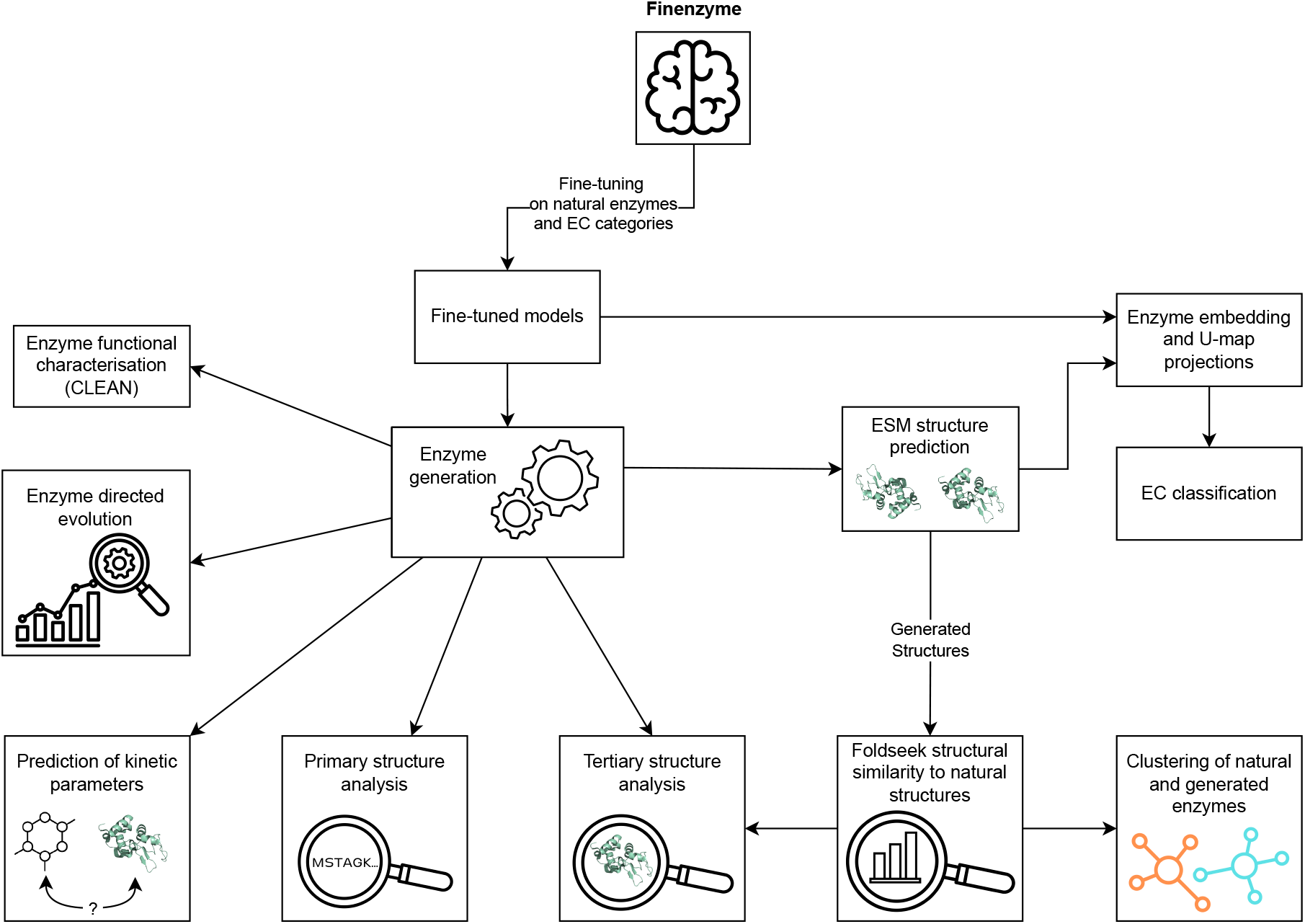
High-level scheme of *Finenzyme* and summary of the main computational tasks performed in this work.

### 2.1 Fine tuning is effective only for low-level EC classes

An important question is whether fine-tuning should be expected to perform equally well across all EC classes or if its effectiveness varies within the EC hierarchy. For instance, high level (more general) classes encompass a broader range of enzymes available for fine-tuning. However, these enzymes are more heterogeneous than those in low level (more specific) classes, raising the question whether the effectiveness of finetuning may depend on the hierarchy of EC categories.

To answer this question, we fine-tuned seven models on general EC classes and seven on more specific EC classes, and we tested their performance at predicting the next amino acid in a sequence. The general EC classes represent broad categories of enzyme functions, such as oxidoreductases (EC1), transferases (EC2), hydrolases (EC 3), lyases (EC 4), isomerases (EC 5), ligases (EC 6), and translocases (EC7). Specific EC classes denote particular enzyme catalysed reactions and in our experiments we used alcohol dehydrogenase (EC 1.1.1.1), DNA methyltransferase (EC 2.1.1.37), cellulase (EC 3.2.1.4), ribulose-bisphosphate carboxylase (EC 4.1.1.39), chorismate mutase (EC 5.4.99.5), biotin ligase (EC 6.3.4.15), and proton-translocating transhydrogenase (EC 7.2.1.1) (see Materials and Methods for details).

For each EC category, we randomly split the data into 90% training and 10% test sets. We also collected a “filtered test set” by selecting test examples having BLAST sequence similarity against the training set less than 70%. We measured the performance of our fine-tuned models at predicting the next amino acid in a sequence in two scenarios : a) Teacher Forcing (TF) in which the LLM is “forced” to predict the next amino acid *x*_*i*_ given the correct previous amino acids, i.e. *x*_*<i*_; b) PreFixed testing (PF) in which the LLM makes a prediction without teacher forcing, using a prefix string for the first *n* = 20 amino acids. In both scenarios, the most probable amino acid is selected at each next amino acid prediction step. We measured performance using three different metrics, namely mean accuracy per-token, mean soft accuracy per-token based on BLOSUM62 (Henikoff and Henikoff, 1992) amino acid substitution matrix, and perplexity (see “Fine-tuning of conditional in Transformers” in Materials and Methods for details).

Fig. 2a compares the teacher forcing results obtained between *Finenzyme* and ProGen models summarizing the distribution of accuracy, soft-accuracy, and perplexity for general and specific EC classes. There is a significant difference in the distributions in all specific EC tests (Wilcoxon signed rank test, *p* − *value* < 10^−4^ for accuracy, soft accuracy and perplexity). Our predictions show that fine-tuned models greatly outperform ProGen when the EC classes are specific (i.e. low in the EC hierarchy), while results are comparable when the EC classes are general. Importantly, the difference in accuracy, soft accuracy and perplexity remains significant also when using the filtered test set (*p* − *value <* 10^−4^), highlighting that *Finenzyme* generalizes better also for out-of-distribution test sets compared to ProGen.

**Figure 2:**
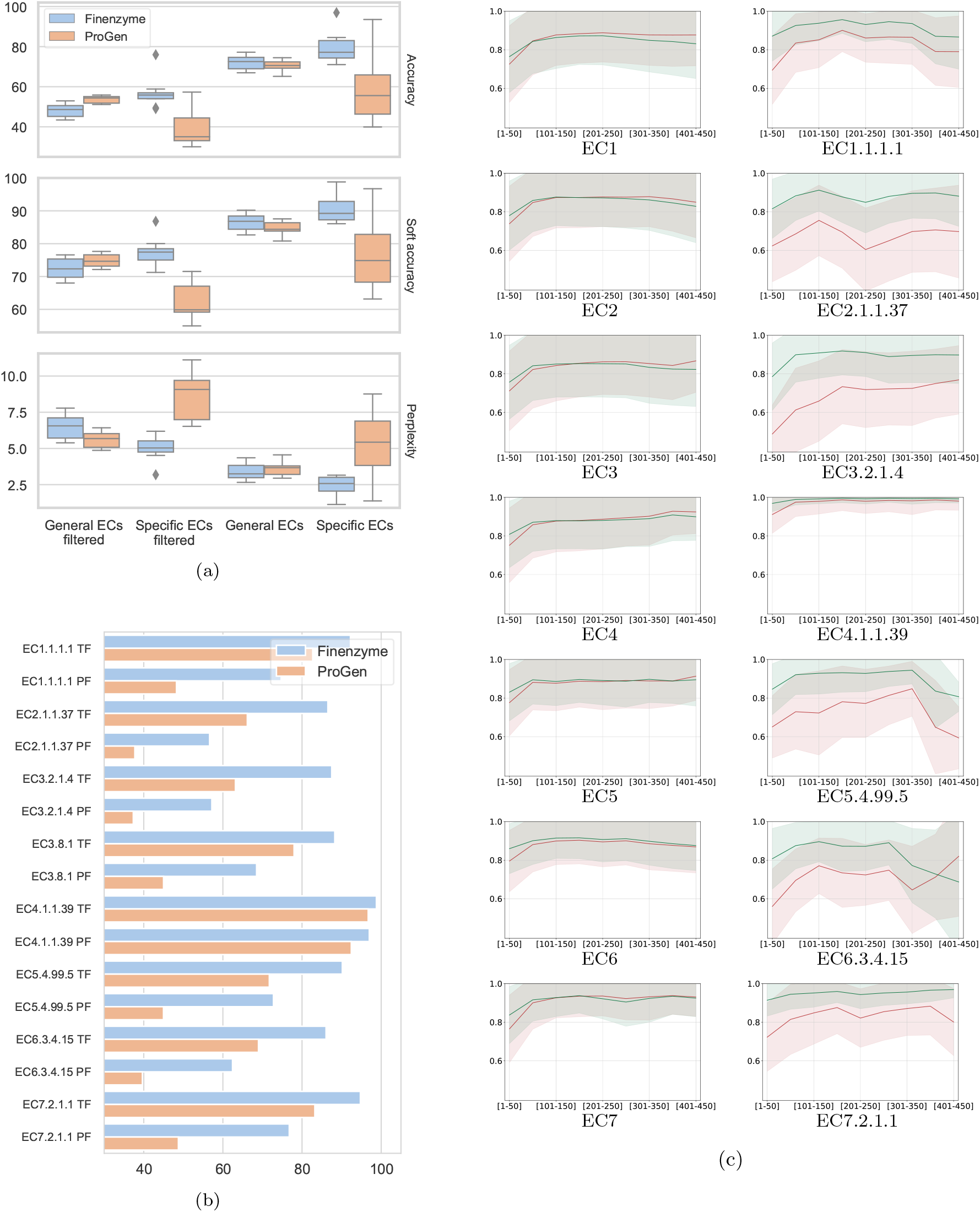
Comparison of ProGen and *Finenzyme* on general and specific EC classes. (a) accuracy (top), soft accuracy (center), perplexity (bottom) compared between general and specific EC classes. “Filtered” denotes whether the test dataset was filtered using BLAST against the training set to retain only sequences with less than 70% identity. (b) soft accuracy of the specific EC classes. “TF” denotes teacher forcing, “PF” testing without teacher forcing with a prefixed chain of 20 amino acids. (c) soft accuracy comparison of *Finenzyme* and ProGen on general (left) and specific (right) EC classes on the full test set. The x-axis reports the position of the amino acid and the y-axis the corresponding average soft-accuracy of the predicted enzymes in ProGen (red line) and *Finenzyme* (green) across the enzymes of the EC category. Shadows represent the standard deviation.

Fig. 2b shows a comparison between TF and PF results using soft-accuracy for the more specific EC classes, confirming that fine-tuned models achieve significantly better results in both scenarios – similar results are also obtained when measuring accuracy and perplexity (Fig. S3, Supplementary Information).

Fig. 2c compares the soft accuracy calculated on general and specific EC categories at each position in the sequence showing that *Finenzyme* consistently outperforms ProGen over the entire sequence length, independently on the amino acid position. Figs. S4– S11 in the Supplementary Information show the same comparison using accuracy, soft accuracy, and perplexity in all test configurations.

### 2.2 *Finenzyme* sequences are similar in sequence, structure and function to natural enzymes

A key aim of this work is to be able to generate enzymes with properties similar to those of the natural ones. Therefore we used *Finenzyme* to generate sequences for the more specific classes and we then carried out experiments to assess their biological plausibility. For each low-level EC category, we generated 2000 enzymes using only the EC keywords. To accomplish this, during fine-tuning we applied top-*p* filtering techniques (Holtzman et al., 2020) for choosing the next generated amino acid, with *p* = 0.75, or *p* = 0.5 and repetition penalty parameter of 1.2. Then we adopted the “global optimal stop-choosing technique” to choose the length of the generated enzyme, (see “Generation of new enzyme sequences” in Material and Methods for details). Below we report our analysis on comparing sequence, structure, and function of the *Finenzyme* sequences to natural enzymes.

### 2.2.1 In depth sequence analysis of the generated enzymes

Fig. 3a shows that the distribution of the length of *Finenzyme* sequences is very similar to the one of the natural enzymes used to train *Finenzyme*. The primary sequences themselves share similarities, as evidenced by the high BLAST max identity scores for all specific EC classes (Fig. 3b). Besides one EC class having all similarities close to 100% (4.1.1.39, ribulose-bisphosphate carboxylase), and one having median similarity slightly below 70% (6.3.4.15, biotin–[biotin carboxyl-carrier protein] ligase), all other EC classes have a median similarity equal to or above 80%.

**Figure 3:**
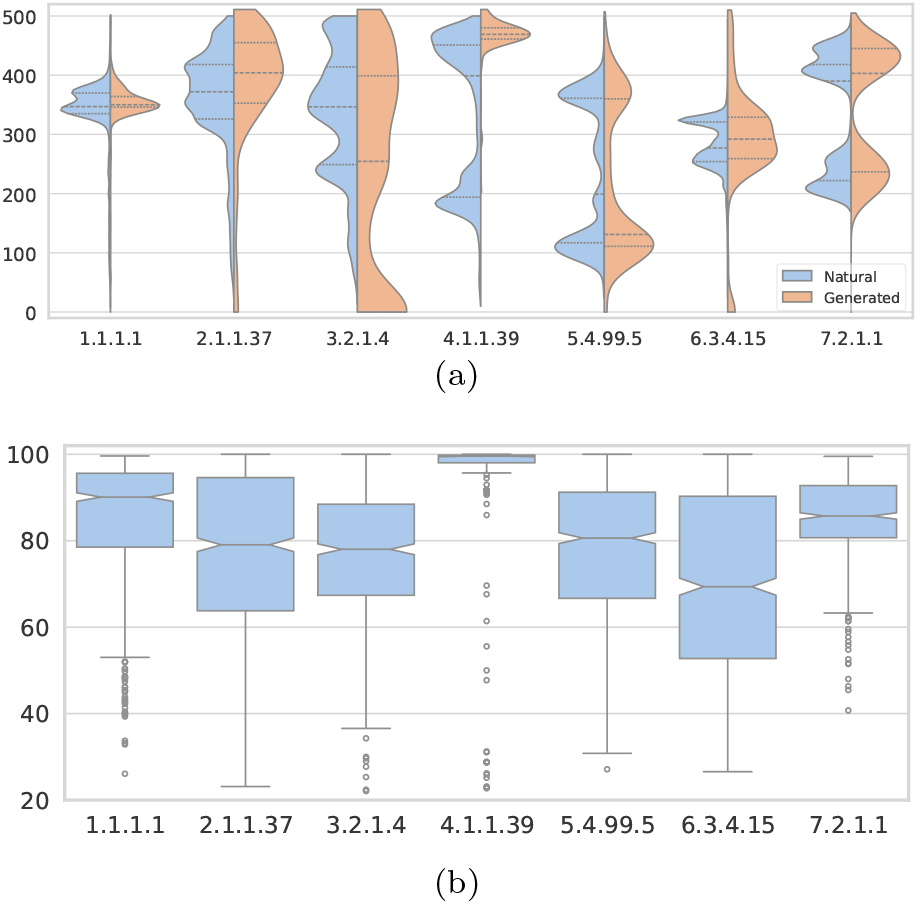
Evaluation of the quality of the primary structure of the generated enzymes. (a) Length distribution of the generated and natural proteins. Violin plots show the length distribution of the natural enzymes included in the training set (blue) and the length distribution of the corresponding generated enzymes (orange). Generation has been performed with top-*p* sampling (*p* = 0.75). (b) Sequence max identity distribution of the generated enzymes. Box plots show the BLAST maximum identity results for each generated protein set with respect to natural proteins of the training set, using only keywords. Generation for (a) and (b) has been performed using top-*p* sampling (*p* = 0.75).

We also observe that a relatively large number of the generated enzymes registers lower primary sequence similarities, also below 40%, showing that *Finenzyme* generates also sequences that diverge from the natural ones.

#### 2.2.2 In depth tertiary structure analysis of the generated enzymes

To assess the tertiary structure similarity between the generated enzymes and the natural ones, we used ESMFold (Lin et al., 2023) to predict structures for *Finenzyme* sequences and applied Foldseek (Van Kempen et al., 2024) to select the most structurally similar pairs of generated and natural enzymes retrieved from PDB according to the Foldseek structural bit scores (see “Evaluation of the quality of *Finenzyme* sequences” in Materials and Methods for details).

Fig.4a shows the relationship between structural similarity (measured by the TM-score) and sequence identity between top-hit Foldseek pairs of generated and natural proteins available in PDB. The TM-score distribution (top x-axis – Fig. 4a) is centered around 0.9, indicating that the structures of the generated and natural proteins are very similar. The distribution of sequence similarity (left y-axis – Fig. 4a) spans a wide range of values, with a mode between 0.3 and 0.4, and has a low correlation with TM-score (Pearson correlation *ρ* = 0.14). *Finenzyme* thus generates primary sequences that differ from those of natural enzymes but may preserve the tertiary structure of the enzymes and thereby probably retain their original enzymatic functions.

**Figure 4:**
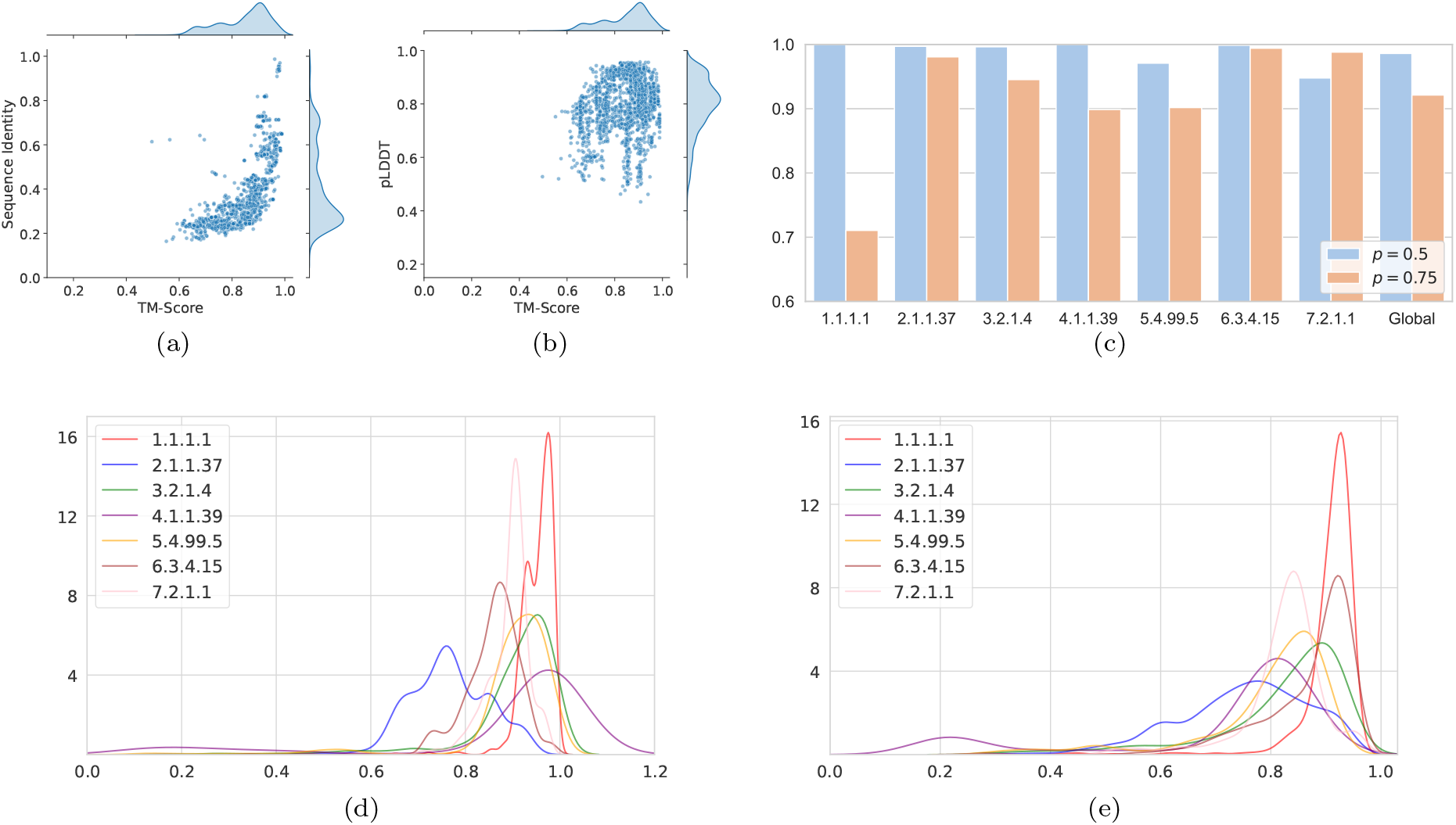
Analysis of the tertiary structure and function of *Finenzyme* generated enzymes. (a) Correlation between structural similarity (TM-score) and sequence identity (BLAST Max ID) between generated and natural enzymes retrieved from PDB across all low-level EC classes. (b) Correlation between ESMFold prediction confidence (pLDDT) and structural similarity to known proteins in the PDB (TM-score). Blue scatterplots (a, b) refer to top-*p* = 0.5 nucleus filtering. (c) Results of CLEAN EC category predictions. Bar plots show the F1 scores for EC number predictions for the generated enzymes, with blue bars representing results of enzymes generated with *p* = 0.50 and orange bars with *p* = 0.75. “Global” represents the weighted average F1 score when considering all EC numbers combined. (d) Distribution of TM-scores, and (e) pLDDT values for each low-level EC category.

Finally, we checked whether the enzyme structural similarities were correlated with the confidence of the ESMfold predictions. Fig. 4b shows that there is indeed a high correlation between the pLDDT values (quantifying the confidence of ESMFold predictions) and TM scores (Pearson correlation *ρ* = 0.64). Details of the TM-score and pLDDT values distribution for each low-level EC category are displayed in Fig. 4d,e. Supplementary Fig. S15 and S16 present the relationships between structural and sequential similarity and ESMFold prediction confidence for each EC class.

#### 2.2.3 *Finenzyme* sequences with low sequence similarity preserve tertiary structure of natural enzymes

We compared predicted structures from *Finenzyme* sequences and the closest match found in the PDB database by Foldseek. Fig. 5 depicts some specific cases of proteins generated by *Finenzyme* having low primary structure similarity but high 3D structural similarity with natural enzymes. More precisely Fig. 5 shows the closest match between the *Finenzyme* sequences (in green) and natural ones (in yellow), considering three different enzymes. Note that in all cases the binding site is predicted with high confidence and provides adequate space for the lig- and (highlighted in violet).

**Figure 5:**
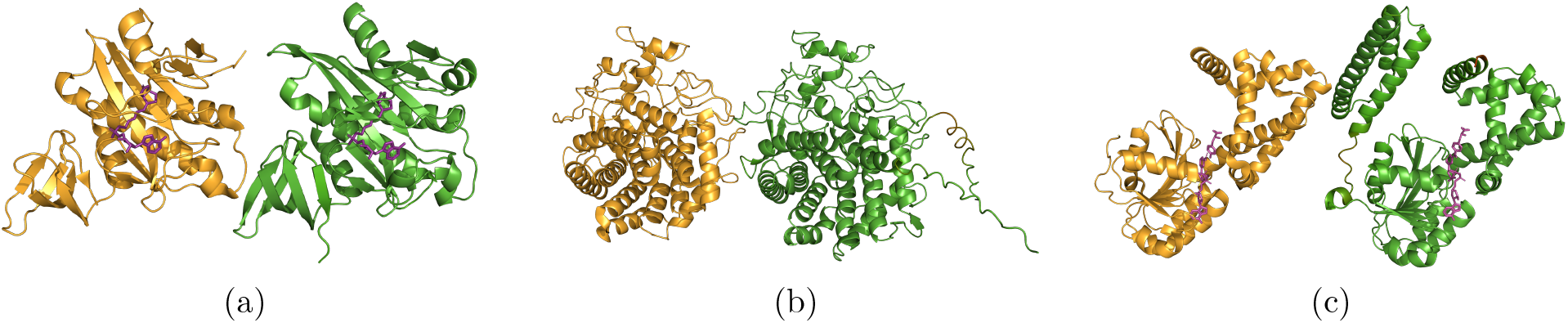
Comparison of the tertiary structure of *Finenzyme*-generated sequences (green) and natural enzymes (yellow). Enzyme generated from (a) EC family 6.3.4.15 (biotin ligase). The target found is “2e41”, a biotin protein ligase (UniProt accession O57883) from *Pyrococcus horikoshii*: sequence similarity = 34.9%, TM-score = 0.94, pLDDT (ESMFold) = 0.95; the binding site of the enzyme is predicted with high confidence and provides adequate space for the ligand (highlighted in violet). (b) EC 3.2.1.4 (cellulase). The PDB target found is “8ihw”, an endoglucanase from *Eisenia fetida*: similarity = 39.6%, TM-score = 0.95, pLDDT (ESMFold) = 0.92. (c) EC 5.4.99.5 (chorismate mutase). The PDB target found is “2pv7”, a prephenate dehydrogenase from *Haemophilus influenzae*: similarity = 49.6%, TM-score = 0.9, pLDDT (ESMFold) = 0.9; the binding site of the enzyme is predicted with high confidence and provides adequate space for the ligand (highlighted in violet).

#### 2.2.4 Functional characterization of the generated enzymes

We predicted the functional properties of *Finenzyme* sequences using CLEAN, a state-of-the-art enzyme function prediction tool (Yu et al., 2023b), that can assign EC numbers to protein sequences (see “EC classification of the generated enzymes” in Materials and Methods for details). The F1 scores for enzymes generated at top-p, with *p* ∈ {0.5, 0.75} are displayed in Fig. 4c (the recall scores can be found in Supplementary Fig. S14). For both generation settings, CLEAN achieves robust predictions with minimal variations across the classes, demonstrating that the functional characteristics are well preserved even when primary sequences diverge from those of natural enzymes.

### 2.3 An in depth computational analysis of hydrolases acting on C-halide compounds (EC 3.8.1)

To demonstrate the possible applications of *Finenzyme*, we turned our attention on an industrially important family of enzymes (EC 3.8.1). Organohalogen compounds are extensively employed across various sectors including agriculture, healthcare, manufacturing, and more, due to their versatility in applications ranging from herbicides and insecticides to refrigerants and synthetic precursors. Despite their broad utility, they represent a significant environmental pollutant (Häggblom and Bossert, 2003), permeating marine and terrestrial ecosystems alike (Gribble and Gribble, 1996). Their chemical stability and resultant persistence in the biosphere poses ongoing environmental and health risks due to the difficulty in degrading these compounds and their toxic effects even at low concentrations (Liu et al., 2019; Shaw, 2010). Enzymes such as dehalogenases have gained considerable interest for their potential in bioremediation and industrial applications, where they transform organohalogens into less harmful compounds (Janssen et al., 2005; Kurihara and Esaki, 2008; Swanson, 1999). Thus, the study and enhancement of dehalogenase activity are pivotal in advancing our capabilities to manage and mitigate the impact of halogenated pollutants globally.

We trained *Finenzyme* with dehalogenases sequences retrieved from Uniprot (The UniProt Consortium, 2023). Fig. S20 shows the number of enzyme sequences belonging to each dehalogenase sub-class. The model was trained by prepending each sequence with the more general 3.8.1 token. Given the low number of sequences for some subclasses, we only prepended more specific tokens (3.8.1 + 3.8.1.2 / 3.8.1 + 3.8.1.5 / 3.8.1 + 3.8.1.3) when more than 1000 instances were available for that specific subclass. We observe that fine-tuning on a set of functionally related EC categories allows for transfer learning and a substantial enlargement of training examples. For instance, with EC 3.8.1.3 we can increase the number of available examples from about 1300 to about 20000, thanks to the addition of the related training examples from EC 3.8.1.2 and 3.8.1.5.

The TF prediction results for each subclass are depicted in Fig. 6a. Compared to ProGen, *Finenzyme* achieved substantial improvement in all the evaluation measures and across all the EC subclasses (Wilcoxon signed rank tests p-value *p <* 10^−5^). Although the improvement is more evident on the most represented subclasses, our results show that this experimental setting boosts performance across all EC subcategories.

**Figure 6:**
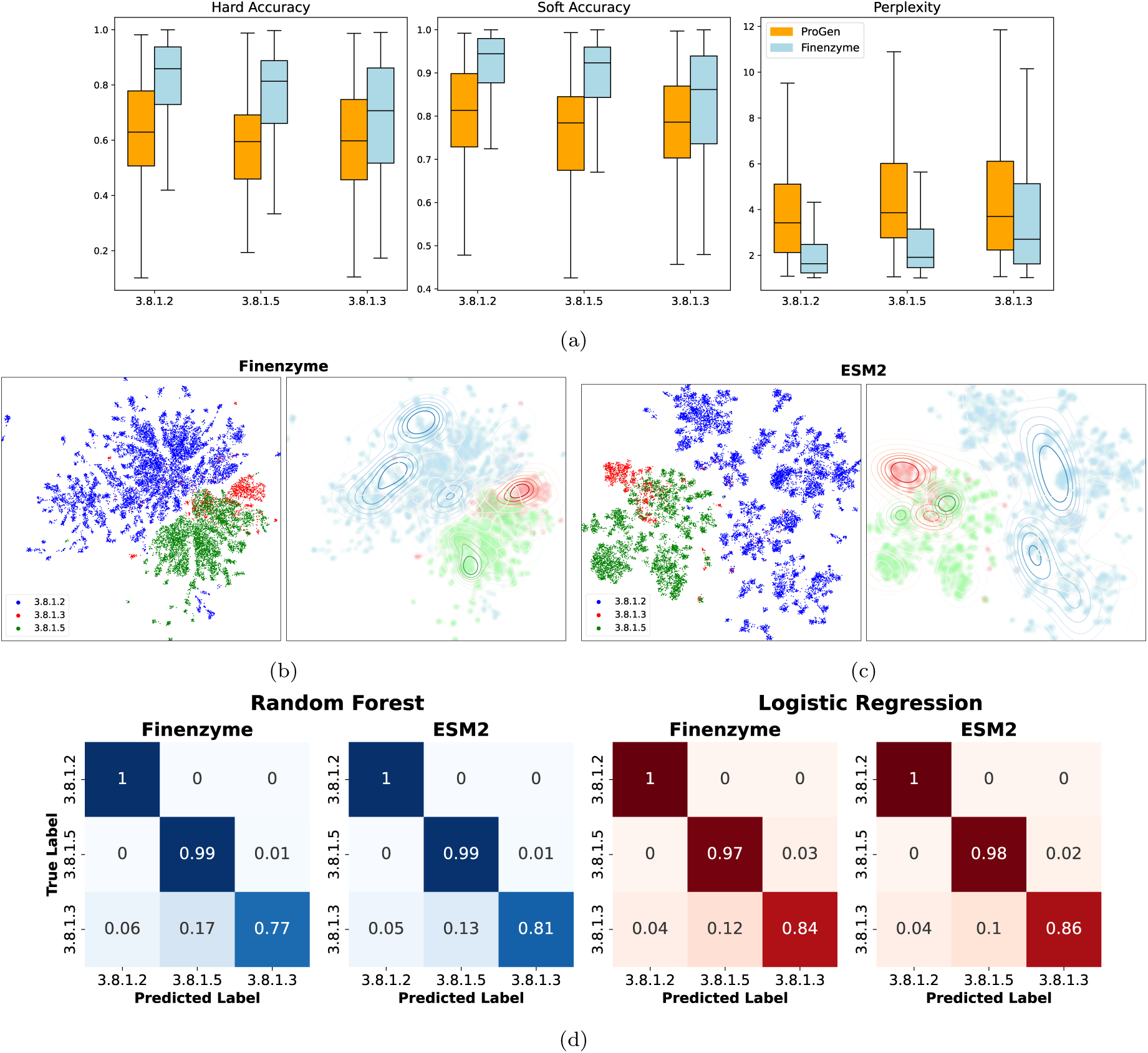
Prediction results and analysis of the embedded representations of natural and *Finenzyme* generated proteins for hydrolases acting on C-halide compounds. (a) Perplexity, hard accuracy, and soft accuracy performance of *Finenzyme* model against Progen for the different dehalogenase subclasses on the test set. (b) UMAP projections of the embedded *Finenzyme* representations of (left) natural enzymes, and (right) generated enzymes. Considering the disparity in cardinality of certain subclasses, we normalized the distribution of the generated sequences through kernel density estimation maps. (c) UMAP projections obtained from the embeddings of ESM2 comparing natural and generated enzymes. (d) Confusion matrices of EC classification tasks performed by Random Forest (blue) and Logistic Regression (red), using the embeddings retrieved from *Finenzyme* and ESM2.

To assess the effectiveness of tokens in influencing model generation toward the correct subclass, we generated 1000 sequences for each subclass using nucleus filtering with top-*p* ∈ {0.5, 0.75}. The lengths of the generated sequences in each experiment, compared to those of the training set, are shown in Fig. S21. Duplicates were removed, and a summary of the generated and natural sequences is provided in Supplementary Table S7. Fig. S22 shows the distribution of the max-ID BLAST sequence identity scores between the generated and natural enzymes across 3.8.1, 3.8.1.2, 3.8.1.3, and 3.8.1.5 EC categories.

#### 2.3.1 Embedded representations of natural and *Finenzyme* generated enzymes are similar

Embeddings extracted from the last PLM layers capture relevant features for protein structure and function (Elnaggar et al., 2022; Rives et al., 2021). Accordingly, we employed embedded representations extracted from the last hidden layer of *Finenzyme* and ESM-2 (Lin et al., 2023) for both natural and *Finenzyme* sequences (details in Section 4.9). Natural sequences with subfamily keywords 3.8.1.2, 3.8.1.5, 3.8.1.3 were used to project the embeddings into 2D representations through UMAP (McInnes et al., 2018). Fig. 6b,c presents the UMAP projections obtained from the embeddings obtained from *Finenzyme* and ESM2, respectively (see “Embedded representations on natural and Finenzyme generated enzymes” in Material and Methods for details). Independently of the method used to generate the embeddings, the UMAP projections of natural and *Finenzyme* embedded representations are very close (Fig. 6b,c, right), indicating that our fine-tuned model captures the main features of the natural enzymes.

The embeddings were then used to train random forest and logistic regression classifiers. Confusion matrices obtained through cross-validation show that both classifiers correctly predict EC subclasses of 3.8.1 using independently *Finenzyme* or ESM2 embeddings (Fig. 6d).

#### 2.3.2 Clustering of the 3D representation of natural and generated enzymes are close

To evaluate the relationship between the structure of the natural and *Finenzyme* sequences, we clustered natural and generated enzymes using Single-Linkage Hierarchical Clustering. The metric used to construct the distance matrix was the E-value retrieved from two experiments that include both the natural and *Finenzyme* sequences: 1) Protein BLAST all-vs-all, which accounts for primary structure similarity; 2) Foldseek all-vs-all, which accounts for both primary and tertiary structure similarity (see “Clustering of natural and *Finenzyme* generated enzymes” in Materials and Methods for details).

In order to provide a more comprehensive database for structural analysis comparison, we removed the constraint for PDB experimentally validated structures, and we predicted the structures for the entire natural dataset using ESMFold. The heatmaps of the adjacency matrices reordered according to Bar-Joseph et al. (2001) clustering (see Section 4.10) are shown in Fig. 7. At the side of each heatmap, kerneldensity estimates allow to visually compare the relative distributions of *Finenzyme* sequences and natural proteins. We can observe that each subclass contains sub-clusters, and clusters of natural and *Finenzyme* enzymes largely overlap.

**Figure 7:**
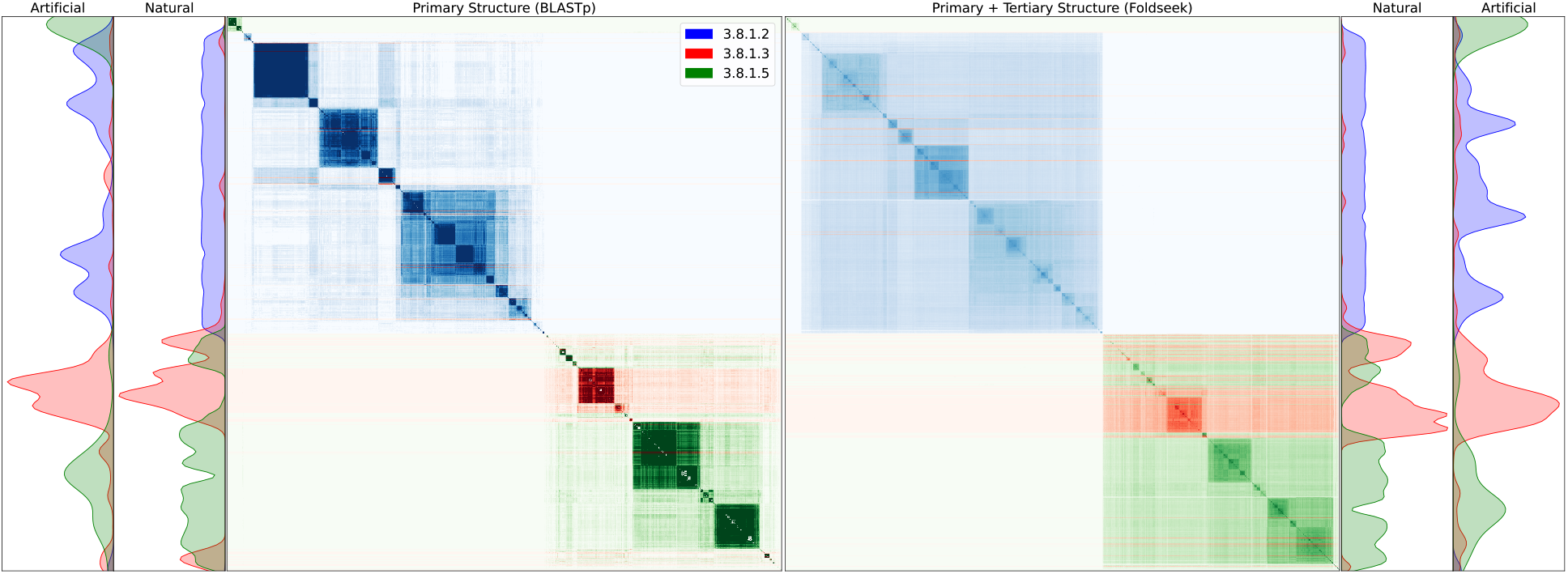
Heatmaps of the clustered natural and *Finenzyme* generated “artificial” enzymes. Different colors highlight different EC categories. Heatmaps are obtained from the adjacency matrix computed through BLAST (left) and Foldseek (right) E-values. On the sides: kernel density estimation of the distribution of natural and *Finenzyme* generated proteins.

#### 2.3.3 *Finenzyme* shows a significantly larger correlation than ProGen between the confidence of its predictions and structural and sequence identity

We compared the confidence in the prediction of *Finenzyme* with ProGen by evaluating the log-likelihood of their generated sequences, which shows how confident a model is in the prediction of an enzyme sequence. Fig. 8a,b shows that *Finenzyme* log-likelihood strongly correlates with pLDDT, TM-score, and sequence identity, and the correlation is significantly larger for *Finenzyme* with respect to ProGen, underscoring that *Finenzyme* is able to predict with higher confidence sequences that are more biologically plausible.

**Figure 8:**
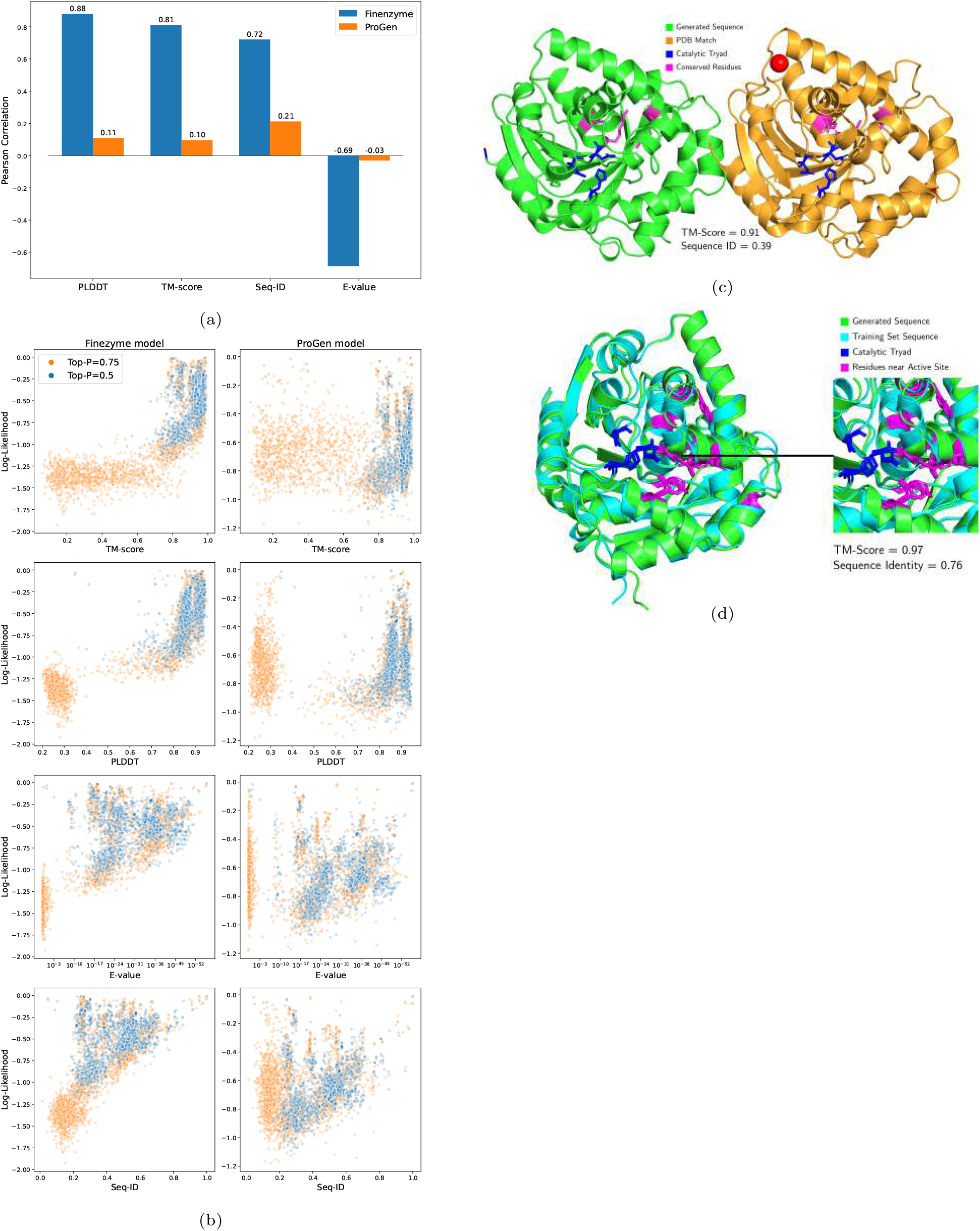
Correlation between the log-likelihood of the predicted enzymes with TM-scores, pLDDT, sequence identity and E-value. (a) Pearson correlation between the log-likelihood scores of *Finenzyme* (blue) and ProGen (orange) against pLDDT, TM-score, sequence identity, and E-value. The metrics are computed either through ESMFold or Foldseek (b) Log-likelihood of the predictions of *Finenzyme* (left) and ProGen (right) as a function of TM-score, pLDDT, E-value and sequence identity. (c) Comparison of the predicted ESMFold 3D structure of the natural enzyme (fluroacetate dehalogenase, in gold) and the corresponding *Finenzyme* protein (green) generated from EC subcategory 3.8.1.3 (fluoracetate dehalogenases). Important residues belonging to the catalytic triad (blue) or favoring the reaction (magenta) are highlighted. (d) The same *Finenzyme* predicted sequence superimposed onto the corresponding natural one retrieved from the AlphaFoldDB. Residues belonging to the catalytic triad are highlighted in blue and others facing the active site in magenta.

Moreover, the fine-tuned model shows a decrease in sequence identity correlation compared to the TM-score, which evaluates the tertiary structure similarity. These results show that *Finenzyme* can better learn the 3D structural plausibility of a sequence rather than its mere primary sequence.

Driven by the ability of our model to preserve structure while altering sequence identity, we compared the predicted structure of one of the enzymes generated by our model using the keyword 3.8.1.3 (fluoroacetate dehalogenases), i.e. the least represented subfamily, with its most similar PDB counterpart. The TM-score between the two structures is 0.91, despite a sequence identity of 0.39 (Fig. 8c). Furthermore, driven by these results, we conducted a literature investigation of the PDB structure. The PDB reference structure (PDB code = 3R3U) corresponds to the fluoroacetate dehalogenase RPA1163, which has been thoroughly studied due to its potential industrial applications (Chan et al., 2011; Kim et al., 2017; Yue et al., 2021).

By examining our predicted structure and comparing it with the real one, we can observe that the catalytic triad has been maintained (Fig. 8c, blue), and several important amino acids necessary for the enzymatic reaction are also well conserved (Fig. 8c, magenta). However some regions of the proteins differ. We hypothesized that it might be due to the generated enzyme resembling the structure of a natural protein for which no PDB structure has been experimentally solved yet. Interestingly, the training set sequence with the highest sequence identity has a predicted structure present in the AlphaFoldDB (Varadi et al., 2024). We, therefore, compared our enzyme with the latter. In this case, the sequence identity reaches 0.76, but the TM-score skyrockets to 0.97 (Fig. 8d). Furthermore, all amino acid residues facing the active site are extremely conserved.

### 2.4 Enzyme directed evolution through *Finenzyme*

The generation of enzymes “evolved” from natural counterparts through *Finenzyme* aims to produce new sequences with well-characterized functional properties.

We focused on three sequences of significant scientific interest: P11766 (alcohol dehydrogenase - EC 1.1.1.1) is characterized by its high activity towards formaldehyde (Engeland et al., 1993); P05102 (EC 2.1.1.37) prevents the incorporation of damaged bases into DNA, maintaining genomic stability and integrity (Mi et al., 1995); Q9SL92 (EC 6.3.4.15) is a potential target for genetic manipulation to improve crop yield and stress tolerance through enhanced metabolic efficiency (Puyaubert et al., 2008).

The generation process, detailed in “Generation of new enzyme sequences” (Material and Methods), used both keywords and amino acid prefix of the targets, yielding a total of 3000 sequences. First, duplicate sequences were filtered out (Supplementary Table S5 provides the count of unique sequences per target). Next, predicted structures were obtained using ESMfold and were then evaluated versus the target enzymes. Structures with a pLDDT greater than 0.7, being the latter a good confidence score, as defined by ESM authors (Rives et al., 2021), were retained, and those with a TM-score to the target greater than 0.7 were selected. Note that these are parameters of the “directed evolution process”, and as such they can be tuned according to specific generation objectives (see S5 for details).

We also matched the generated enzymes against the Protein Data Bank (PDB) using Foldseek. Supplementary Table S5 shows that for each target enzyme we obtained a relatively large set of novel enzymes that satisfy the above criteria. Subsequently, we clustered the filtered sequences using the Foldseek representation of enzymes and the MMseqs suite for fast and deep clustering adapted to the “3Di” alphabet of Foldseek (Hauser et al., 2016). From each cluster we extracted a representative enzyme and performed multiple sequence alignments based on the Foldseek enzyme representations. The alignment results relative to P05102 are visualized using a circular tree (Fig. 9a), that includes the natural enzyme targets and the corresponding enzymes selected through directed evolution. Analogous circular tree figures for P11766 and Q9SL92 are available in the Supplementary Information (Fig. S17, and S18).

**Figure 9:**
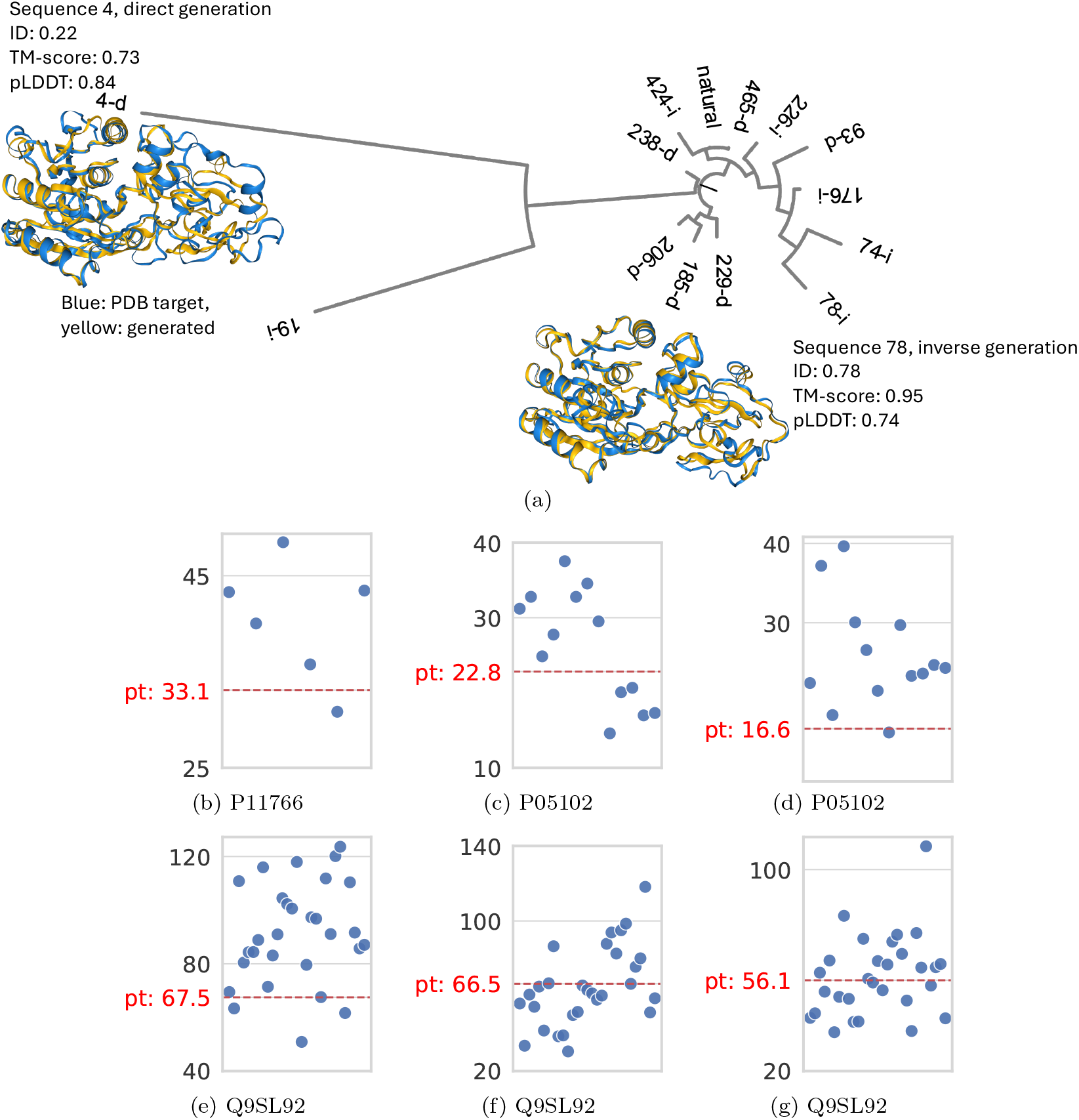
*Finenzyme* directed evolution and evaluation of the kinetic parameters. (a) Circular tree representing the directed evolution of enzyme P05102 (PDB structure identifier 1fjx) (b–g) Kinetic parameter *k*_m_ prediction using UniKP. Each scatter plot displays the predicted *k*_m_ between the natural sequence and the substrate as a red line; the blue points are predictions of *Finenzyme* proteins generated through directed evolution. The enzyme-substrate pairs are (b) P11766 and 20-HETE, (c) P05102 and DNA, (d) P05102 and S-adenosyl-L-methionine, (e) Q9SL92 and biotin, (f) Q9SL92 and ATP, (g) Q9SL92 and methylcrotonoyl-CoA carboxylase. Values are displayed in *µ*M.

Finally, Michaelis-Menten kinetic parameters of the natural enzyme target and of the enzymes selected by directed evolution with *Finenzyme* were predicted using UniKP (Yu et al. (2023a) (see “Prediction of the kinetic parameters of the generated enzymes” in Material and Methods). The results of these predictions are depicted in Fig. 9b–g, highlighting the functional capabilities of both natural and evolved enzymes. Supplementary Table S6 describes the substrate/protein interaction data available in UniProt for the chosen targets, and the predicted *k*_*m*_ values for the true targets.

## 3 Discussion

Several studies demonstrated that fine-tuning can improve PLM performance in various tasks and settings (Sledzieski et al., 2023; Wang et al., 2024b; El-naggar et al., 2022); however, other works highlighted that this technique does not necessarily lead to better results (Tinn et al., 2023; Schmirler et al., 2024).

In the context of enzymes, we showed under which conditions fine-tuning a conditional Transformer can lead to significantly better results compared to the “general” PLM. Fine-tuning specific EC categories and exploiting specific keywords in the conditional learning process boosts performance. On the other hand, for more general categories, e.g., the top highlevel EC categories, this strategy does not yield better results (Fig. 3c). This can be explained by the fact that general models like ProGen can generalize on overly represented EC categories, by leveraging their knowledge about the large corpus of data on which they have been trained. Conversely, the general model cannot leverage specific knowledge to accurately predict underrepresented and functionally characterized EC categories, while fine-tuned models can learn and focus on specific and well-characterized EC categories.

Analysis of *Finenzyme* sequences using ESMFold, Foldseek, and other state-of-the-art software tools, reveals that the tertiary structure is highly conserved compared to natural enzymes (average TM-score ≃ 0.9), while the sequence identity is less preserved (Fig. 4). This indicates that *Finenzyme* sequences can diverge from natural ones but tend to preserve their 3D structure. The in silico prediction of EC categories of the generated and natural enzymes confirms that *Finenzyme* generated enzymes maintain the same function of the natural ones (Fig. 4c). Moreover, through *Finenzyme* directed evolution, we showed that the Michaelis-Menten kinetic parameters of *Finenzyme* sequences are comparable to, and sometimes even enhance, those of the corresponding natural enzymes (Fig. 9). This opens the way for possible *Finenzyme* industrial or medical applications.

For overly underrepresented EC categories, we propose transfer-learning techniques by jointly finetuning the conditional Transformer on functionally related EC categories while maintaining their specificity using their corresponding keywords. Results with EC 3.8.1 low-level categories show that *Finenzyme* can simultaneously learn multiple functionally related EC categories, using multiple keywords and improving the cardinality of the training examples.

Embedded representations of natural and generated enzymes obtained from the last hidden layer of both *Finenzyme* and ESMFold models span a similar distribution (Fig. 6b,c), and the embedded representations themselves can be used to correctly classify the EC categories (Fig. 6d).

Hierarchical clustering of the 3D structures of both *Finenzyme* generated sequences and the corresponding natural ones yields largely superposed clusters, and every EC category includes sub-clusters (Fig. 7). This is expected since EC numbers do not specify enzymes but rather enzyme-catalyzed reactions. Namely, if different enzymes catalyze the same reaction, they receive the same EC number, regardless of their structural similarity (Bansal et al., 2022). Moreover, through convergent evolution, completely different protein folds can catalyze an identical reaction and therefore would be assigned the same EC number (Omelchenko et al., 2010). However, the smoothness of the Foldseek heatmap (Fig. 7) shows that tertiary structures are more maintained among clusters than primary sequences, confirming the relationships we showed between TM-score and sequence identity scores (Fig. 4).

The intra-cluster structure depicted in Fig. 7 suggests to apply fine-tuning to generate specific protein domains, as proposed by Nijkamp et al. (2023). Another solution could be to cluster the dataset before training and assign additional sub-keywords to each sub-cluster. This can be accomplished by fine-tuning *Finenzyme* on the discovered sub-clusters and using multiple keywords of other functionally similar enzyme categories to overcome the underrepresentation of small enzyme subcategories.

Nevertheless, by looking at the kernel density distributions on the sides of the heatmaps, we found that *Finenzyme* sequences span all the main natural clusters fairly well (Fig. 7).

The log-likelihood of the *Finenzyme* predictions are significantly more correlated than ProGen with the TM-score, pLDDT, E-value and sequence identity, showing that fine-tuning can significantly boost the reliability and the confidence of the predicted sequences (Fig. 8a,b). Analysis of the tertiary structure of generated and natural enzymes not only outlines that the overall 3D structure is preserved but also that catalytic sites are highly conserved (Fig. 8c,d).

Finally, on the basis of the reliability and biological plausibility of *Finenzyme* sequences, we propose a “*Finenzyme* driven directed evolution” process that can be applied to direct the synthesis of enzymes with structure and function similar to a specific target (Fig. 9), thus simplifying the in silico synthesis of functionally characterized enzymes.

A limitation of this work is represented by the relatively low number of 3D enzyme structures available in PDB. Indeed, to assess the structural similarity between *Finenzyme* generated proteins and natural ones, we compared the ESM2 estimated 3D structure of generated enzymes with the experimentally validated 3D structures of the enzymes retrieved in PDB. If on the one hand this results in a comparison with experimentally verified structures, on the other hand we need to outline that PDB stores only a relatively limited subset of 3D representations of enzymes. This in turn can explain why the sequence similarity is on the average lower when we compare *Finenzyme* sequences versus PDB enzymes (Fig. 4a) than versus enzymes sequences used to train *Finenzyme* (Fig. 3b). However, in both cases the structural similarity, measured by the TM-score, is high (on the average larger than 0.9), witnessing that *Finenzyme* can effectively learn the structural and functional features of the natural enzymes, independently of the corresponding sequence similarity.

Another related limitation of this work is that we needed to estimate the tertiary structure of *Finenzyme* sequences with computational models (ESM2). Even if these methods showed excellent results for the prediction of the protein structure from sequence, we cannot exclude that in some cases we can obtain relatively incorrect predictions.

Overall, our experimental results and analyses support that *Finenzyme* fine-tuning of conditional Transformers constitutes a reliable first step for enzyme engineering to overcome the vast combinatorial space of possible targeted mutations. Producing enzymes with low sequence identity, but high structural similarity through a relatively computationally inexpensive PLM fine-tuning procedure can dramatically boost and assist targeted enzyme engineering tasks.

## 4 Materials and methods

### 4.1 Enzyme Commission taxonomy

The Enzyme Commission (EC) taxonomy (Webb et al., 1992) is a classification system for enzymes, based on the chemical reactions they catalyze. This system categorizes enzymes into four numerical levels, forming a unique EC number for each type of enzyme-catalyzed reaction. For instance, the enzyme class 3.2.1.4 refers to a cellulase, i.e. enzymes that catalyze the reaction of endohydrolysis of (1-4)-beta-D-glucosidic linkages in cellulose, lichenin, and cereal beta-D-glucans. The first number, 3. refers to the general class of the catalyzed reactions: in this case hydrolases that catalyze the hydrolytic cleavage of C-O, C-N, C-C, and some other bonds. The second number (3.2) refers to enzymes that hydrolyze glycosyl compounds. The third number (3.2.1, glycosidases) focuses on specific glycosil compounds on which the enzyme acts, i.e. hydrolyzing O- and S-glycosyl compounds and, finally, 3.2.1.4 refers to cellulases.

### 4.2 Enzyme datasets

In our experiments, we considered both top-level general EC classes and low-level specific EC classes. The top-level classes represent broad categories of enzyme functions, such as oxidoreductases (EC1), transferases (EC2), hydrolases (EC 3), lyases (EC 4), isomerases (EC 5), ligases (EC 6), and translocases (EC7). Each top-level class is further divided into subclasses and sub-subclasses, leading to specific EC numbers that denote particular enzyme-catalyzed reactions. We considered various low-level EC classes, namely, alcohol dehydrogenase (EC 1.1.1.1), DNA methyltransferase (EC 2.1.1.37), hydrolases acting on c-halide compounds (EC 3.8.1), cellulase (EC 3.2.1.4), ribulose-bisphosphate carboxylase (EC 4.1.1.39), chorismate mutase (EC 5.4.99.5), biotin ligase (EC 6.3.4.15), and proton-translocating transhydrogenase (EC 7.2.1.1).

Data sources for our prediction tasks include enzyme sequence datasets downloaded from UniProt (The UniProt Consortium, 2023). The considered sequences were filtered to include only those with length between 10 and 500, and duplicates were removed to avoid redundancy. Top-level classes were further filtered to maintain reviewed-only sequences, i.e., high-quality UniProtKB/Swiss-Prot enzymes curated by experts including non-redundant protein sequence records. This filter was not applied to the most specific EC categories, due to their small cardinality. Hence, to achieve a feasible number of examples for fine-tuning LLMs, non-reviewed sequences have been added to low-level EC categories, i.e., UniProtKB/TrEMBL records that include highquality enzymes enriched with automatic annotation and classification. Supplementary Section S1 and Table S1 in Supplementary Information summarize the main characteristics of the EC categories and of the enzyme datasets included in our experiments.

### 4.3 Conditional Transformers for protein learning and generation

In our experiments, we used a pre-trained model, i.e. ProGen (Madani et al., 2023), that adopted the CTRL conditional Transformer architecture (Keskar et al., 2019), which employs keywords to guide the generation of texts. ProGen generates an amino acid sequence in an autoregressive way, and moreover leverages prepending functional tags to guide the generation. These functional tags can denote a protein family, a Gene Ontology term, or other properties of the protein.

More precisely, given a training instance, represented by a protein sequence *x* = {*x*_1_, *x*_2_, …, *x*_*m*_} and its related keywords *t*, where *x*_*i*_, 1 ≤ *i* ≤ *m*, represents the amino acids, the model learns through back-propagation the conditioned probability *p*(*x*|*t*), that can be factorized using the chain rule of probability:

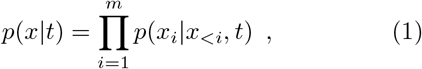

where *p*(*x*_*i*_|*x*_*<i*_, *t*) denotes the conditional probability of *x*_*i*_ given all preceding elements *x*_1_, …, *x*_*i*−1_ and the functional tags *t*.

This formulation breaks down protein language modeling into a next-amino acid prediction task. Consequently, the pre-trained model with parameters *θ* can be trained to minimize the negative log-likelihood over a dataset of sequences *D* = {(*t, x*)^*k*=1^, …, (*t, x*)^*k*=|*D*|^}:

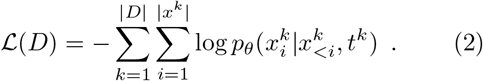

Thus, by acquiring knowledge about the conditional probability distribution, the protein language model can generate new sequences 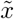 of length *m* by sequentially sampling its components: 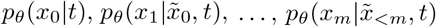

The ProGen architecture features an internal embedding dimension of *d* = 1280, and an inner dimension of *f* = 8192, 36 layers, and 16 attention heads per layer. Each layer includes dropout with a probability of 0.1 following the residual connections, and token embeddings are tied to the final output layer embeddings.

### 4.4 Fine-tuning of conditional Transformers

The goal of fine-tuning is to leverage the knowledge acquired from millions of protein sequences and transfer the “learned knowledge” encapsulated in the weights of the pre-trained ProGen model into a new model fine-tuned to learn the language of a specific EC category of enzymes.

We specialized the models to learn the enzyme characteristics of specific EC classes through a dual approach: a) Conditional decoder learning: generating enzymes conditioned to a specific EC tag; b) Transfer learning and fine-tuning of a pre-trained model on specific EC classes.

For conditioning the models to learn and generate functionally characterized enzymes we prefixed each enzyme sequence with a conditional tag representing the EC class to which the enzyme belongs to. The new keywords for the new EC classes not included in the pre-trained ProGen model have been encoded and added to the fine-tuned models (see Supplementary Section S2 for details about the coding of the EC categories).

The prefixed tags explicitly provide functional information and exert control over the generation process, thus constraining enzymes characterized by specific properties. To allow the models to learn and generate sequences even without an initial keyword, during training we randomly removed the initial keyword with a probability equal to 0.2.

We fine-tuned seven models specialized on the general EC classes and seven on the specific EC classes (see “Enzyme Commission taxonomy” in Materials and Methods for details), according to the experimental set-up described in “Enzyme datasets”.

For each EC category, we randomly split the data into 90% training and 10% test sets. In addition, for each EC category two test sets were prepared: 1) a full test set that includes all available test data; and b) a filtered test set, with examples having BLAST sequence similarity against the training set less than 70%. We further defined two prediction tasks: a) Teacher forcing (TF) testing: next amino-acid prediction is performed by “forcing” the LLM to predict the next amino acid *x*_*i*_ using the correct *x*_*<i*_ and not the currently predicted 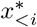 previous sequence, where *x*_*<i*_ represents the enzyme sequence {*x*_1_, *x*_2_, …, *x*_*i*−1_} that precedes the amino acid *x*_*i*_, and 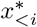 the predicted sequence before the amino acid *x*_i_. b) Prefixed testing (PF): prediction using a prefix string for the *n* = 20 amino acids without teacher forcing. This time the model predicts the next amino acid *x*_*i*_ using 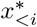, but starts enjoying the correct prefix of *n* = 20 amino acids. In other words, while PF uses the previous predicted sequence to predict the next amino acid, TF uses the correct “true” sequence instead.

During the prediction, top-*k* sampling with *k* = 1 was employed, i.e. the most probable next amino acid is selected at each prediction step. The metrics computed during the prediction tasks are mean hard accuracy per-token, mean soft accuracy per-token based on BLOSUM62 (Henikoff and Henikoff, 1992) amino acid substitution matrix, and perplexity, that is the exponent of the cross-entropy loss calculated over each token in the dataset. Lower perplexity indicates a higher-quality model.

During the learning process, proteins are invariant to the direction of sequence generation; thus we reversed the amino acid sequence with a probability equal to 20%. All models were trained on all weights for about four epochs^1^. The learning rate was set at 0.0001, and the batch size of 2 was used, representing the number of training examples in one iteration. Additionally, a warm-up period of 1000 iterations was implemented, during which the learning rate was gradually increased. To prevent the “exploding gradient” issue common in deep neural networks, gradient norm clipping with a norm of 0.25 was applied. The Adam (Adaptive Moment Estimation) algorithm (Kingma and Ba, 2014) was utilized to compute adaptive learning rates for each parameter, ensuring efficient optimization.

For training and testing the models, we used two multi-processor servers equipped with 128 GB of RAM and NVIDIA *A*100 GPU accelerators.

### 4.5 Generation of new enzyme sequences

Fine-tuned *Finenzyme* models can be used to generate new enzymes by using one of the following two strategies (detailed in the following): (a) Enzyme generation by scratch using keywords only; (b) Enzyme generation from a prefixed sequence of amino acids and keywords.

#### (a) Enzyme generation with keywords only

We started to generate new enzymes using a prompt with keywords only. We applied top-*p* filtering techniques (Holtzman et al., 2020) for choosing the generated amino acid, with *p* ∈ {0.5, 0.75} and we fixed a repetition penalty parameter equal to 1.2. We applied top-*p* nucleus filtering and randomly sampled from this filtered distribution, to let the model explore the most probable amino acid combinations. When the token distribution is peaked, the nucleus will contain fewer tokens with significantly higher probabilities. On the other hand, in flatter distributions, random sampling of the nucleus allows the model to enhance the diversity of the generated enzymes.

We introduced the keyword ‘stop’ learned during fine-tuning to save sequences shorter than 512 amino acids. To choose the length of the generated enzyme, we adopted the “global optimal stop-choosing technique”, i.e. for each position of the sequence we stored the predicted probability of the stop token and then selected the position with the highest probability to set the enzyme final length.

#### (b) Enzyme generation from a keyword plus prefixed sequence of amino acids

Beside the keyword, the second approach for novel sequence generation used a prefixed sequence as a prompt by growing an input ranging from 1 to 250 amino acids (two times per input size, hence generating 500 proteins in total) with *p* = 0.75 for each natural enzyme used as prefixed sequence. We then repeated the process using the flipped sequences.

During generation, a repetition penalty over the last four amino acids was implemented to address the well-known problem of repetitions of the last predicted tokens.

### 4.6 Evaluation of the quality of *Finenzyme* sequences

For each specific EC number, we generated 1000 proteins using top-*p* filtering with *p* ∈ {0.5, 0.75}, resulting in a total of 14000 sequences generated from scratch using only the functional tag as a prompt. After filtering for duplicate sequences, we computationally analyzed the resulting 6885 unique identified enzymes (Supplementary Table S4 presents the distribution of duplicated sequences). We analyzed their similarity with natural proteins considering both their primary and tertiary structure.

For the primary sequence, we evaluated the BLASTp max identity score of *Finenzyme* generated sequences versus natural sequences available both in the EC training set and in PDB, considering the full length of the enzymes, and using BLASTp default parameters. We also compared the length distribution of the generated proteins with respect to natural ones.

To evaluate the 3D quality of the generated enzymes, their structures were predicted with ESM-Fold (Lin et al., 2023) and then compared with those of natural proteins available in the PDB database. The structural similarity was evaluated using the Template Similarity score (TM) (Zhang and Skolnick, 2004).

To identify the natural protein most similar to each generated enzyme we used the Foldseek top-hit for each generated enzyme vs. the PDB database, by using a “structural bit” score, computed as the product of the bit score from the Smith-Waterman algorithm and the geometric mean of the average Local Distance Difference Test (LDDT) and TM scores (Van Kempen et al., 2024).

### 4.7 EC classification of the generated enzymes

To assess the EC classification of the generated enzymes, we used CLEAN (Yu et al., 2023b), a state-of-the-art tool designed to predict enzyme functions accurately. CLEAN leverages embeddings from the ESM-1b model (Rives et al., 2021) to create detailed vector representations of amino acid sequences, which effectively capture the functional similarities between enzymes. These embeddings are processed using a neural network trained with a contrastive loss function, designed to cluster embeddings of sequences with the same EC number while pushing sequences belonging to different EC classes apart.

During the prediction phase, the query enzyme embedding is compared to the centroids of EC number clusters previously established during the training phase. EC numbers are assigned based on the significant proximity of the query’s embedding to these centroids. Two assignment methods are used: the maximum separation method, which selects EC numbers whose centroids are most distinctly separate from others, and the p-value-based method, which identifies EC numbers with statistical significance compared to a background distribution. Since the two methods exhibit similar prediction performances, Fig 4c shows only the results using the maximum separation method.

### 4.8 Prediction of the kinetic parameters of the generated enzymes

To assess the catalytic quality of *Finenzyme* sequences and substrates, we used UniKP (Yu et al., 2023a), a tool designed to predict the Michaelis-Menten kinetic parameters *k*_*cat*_, *K*_*m*_, and *k*_*cat*_*/K*_*m*_. UniKP leverages machine learning models and protein representation techniques. It employs the ProtT5 (Elnaggar et al., 2022) model to encode enzyme sequences and a SMILES Transformer (Honda et al., 2019) to represent substrate structures, creating detailed vector representations of both. These representations are then processed by an Extra Tree ensemble model (Geurts et al., 2006), which predicts the kinetic parameters.

### 4.9 Embedded representations on natural and *Finenzyme* generated enzymes

Both natural and generated enzymes embeddings were retrieved from the last hidden layer of *Finenzyme* and ESM-2 (Lin et al., 2023). As the last hidden layer of *Finenzyme* has a dimension 512 *×* 1280, and the ESM-2 last hidden layer size is 1024 × 5120, average pooling was applied to turn the embeddings into vectors of size 1280 and 5120 respectively. Padding tokens were masked during the computation.

Then, the embeddings were used to learn a 2D latent space utilizing UMAP (Uniform Manifold Approximation and Projection for Dimension Reduction) (McInnes et al., 2018), i.e. the embedded representations of the enzymes were projected into a bidimensional latent space^2^. The embeddings of the full dataset were also used to train a random forest classifier for predicting EC classes^3^. K-fold crossvalidation with *k* = 5 was applied to evaluate the generalization performance.

Using the same experimental setting we trained a logistic regression classifier^4^ with l-BFGS optimizer (Liu and Nocedal, 1989). Considering the size of the embeddings compared to the low cardinality of the data, an “L2 penalty” was applied.

### 4.10 Clustering of natural and *Finenzyme* generated enzymes

We clustered both natural and *Finenzyme* sequences using single-linkage hierarchical clustering^5^. Prior to clustering, we filtered out sequences from natural and generated ones that did not have any subfamily keyword assigned (only 3.8.1.-). To compute the similarities among all sequences, we utilized two algorithms. Firstly, BLASTp (Altschul et al., 1997), which accounts only for sequence similarity; secondly, Foldseek (Van Kempen et al., 2024), which considers both primary sequence and tertiary structure. The algorithms were both run in an “all-vs-all” manner. Finally, the E-value for both algorithms was used to build the distance matrix 𝒟 similarly to the work of Krause et al. (2005). All sequence pairs whose E-value was worse than 0.05 or with no hits were assumed to be unrelated and their distance was set to 1. The linkage matrix computed by hierarchical clustering is reordered to minimize the distance between successive leaves (Bar-Joseph et al., 2001). This reordering is also applied to the distance matrix 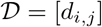 and results in a reordered adjacency matrix 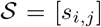 whose heatmap provides a more intuitive and visually coherent representation of the clustering structure. The adjacency matrix 𝒮 is derived from the distance matrix 𝒟 as follows:

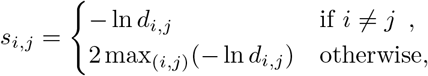

where the self-similarities (on the diagonal) were set to twice the maximum of non-diagonal entries to have large enough values, similar to the approach proposed by Krause et al. (2005).

## 5 Code and data availability

The *Finenzyme* code, the scripts to reproduce the experiments and tutorials to fine-tune *Finenzyme* on any EC category are available from https://github.com/AnacletoLAB/Finenzyme.

The datasets used to train *Finenzyme* were downloaded from UniProt. Details are available from the notebook tutorial in the GitHub repository.

## 7 Author contributions statement

M.N., E.S., G.V., and E.C. designed the overall structure of this work and the experiments. M.N. and

E.S. implemented the code of *Finenzyme*. M.N., E.S., R.J. and A.G. executed the experiments. M.N., E.S., M.M., Em.C. analyzed the results. G.V., E.C., P.R. and A.P. supervised the overall work. G.V., E.C. and A.P. drafted the paper, all the authors revised and approved the final manuscript.

## 8 Acknowledgments

This work was supported by National Center for Gene Therapy and Drugs Based on RNA Technology—MUR (Project no. CN 00000041) funded by NextGeneration EU program.

## Footnotes

1 The number of fine-tuning epochs has been selected according to early-stopping strategies.

2 The implementation of the algorithm was retrieved from the Python package `umap`. The distance metric used was the cosine similarity, with parameters `min` `dist` = 1, and `n` `neighbors` = 100. These settings were kept equal for the embeddings retrieved from both models.

3 We applied the `RandomForestClassifier` from the `sklearn` Python package. The number of trees was set to 2000 while all the other hyperparameters were set to their default values.

4 We applied the `LogisticRegression` implementation from the Python package `sklearn`

5 Single-linkage hierarchical clustering was performed by using `hierarchy.linkage` implementation from the `scipy` library.

