## Supplementary Information for "Fine-tuning of conditional Transformers for the generation of functionally characterized enzymes"

August 10, 2024

#### Contents

**S1 Enzyme datasets used in the experiments** **2**

**S2 Model architecture and keywords encoding** **3**

**S3 Test results** **4**

**S4 Only-keywords generation, CLEAN results, and per-class analysis of ESMFold predictions** **17**

**S5 Prefixed generation and directed evolution** **20**

**S6 *Finenzyme* generation of dehalogenases (EC 3.8.1)** **22**

### S1 Enzyme datasets used in the experiments

The datasets used in our experiments have been downloaded from the UniProt database, specifically focusing on Enzyme Commission (EC) numbers. To create the datasets, we initially collected all available sequences for each EC number. The sequences were then filtered to remove those that were shorter than 10 amino acids (partial reads) or longer than 500 amino acids. This step ensures that the sequences used in our analysis fall within the model maximum input size. After filtering, duplicate sequences were identified and removed to ensure that each sequence in the dataset was unique. Table S1 summarizes the datasets used in the experiments.

In addition to the main datasets, that have been divided in test (10%) and training sets (90%), we generated reduced test sets starting from the full test set for further validation. These test sets consist of sequences with low similarity compared to the training set to ensure a diverse and representative sample. Filtering involved maintaining sequences with less than 70% sequence similarity computed through BLAST algorithm between the full test set and the training set. Table S2 summarizes these reduced test sets.

| EC Number | Initial Seq. | Filtered <10 | Filtered >500 | Duplicates | Resulting Seq. | Mean Length | Std Length |
|---|---|---|---|---|---|---|---|
| 1 | 35286 | 19 | 7067 | 4092 | 24127 | 328.167 | 94.283 |
| 2 | 97889 | 10 | 18820 | 13693 | 65366 | 311.707 | 93.823 |
| 3 | 63560 | 16 | 15246 | 8199 | 40099 | 279.838 | 110.236 |
| 4 | 26479 | 5 | 4015 | 4000 | 18459 | 300.492 | 108.269 |
| 5 | 16238 | 1 | 3483 | 2336 | 10418 | 312.446 | 101.39 |
| 6 | 29297 | 0 | 12389 | 2732 | 14176 | 383.034 | 93.436 |
| 7 | 14720 | 0 | 4373 | 1759 | 8588 | 277.884 | 121.996 |
| 1.1.1.1 | 52818 | 1 | 922 | 5236 | 46660 | 335.44 | 64.73 |
| 2.1.1.37 | 58050 | 0 | 16568 | 3878 | 37604 | 359.91 | 86.16 |
| 3.2.1.4 | 46042 | 2 | 15492 | 1780 | 28768 | 334.24 | 106.05 |
| 4.1.1.39 | 173972 | 3 | 260 | 47827 | 126143 | 334.83 | 127.23 |
| 5.4.99.5 | 41840 | 0 | 197 | 5679 | 35964 | 233.08 | 119.35 |
| 6.3.4.15 | 36935 | 0 | 213 | 2721 | 34001 | 282.24 | 42.16 |
| 7.2.1.1 | 39676 | 0 | 168 | 5589 | 33919 | 331.26 | 99.7 |

Table S1: Datasets summary for families with different EC numbers.

| EC Number | Initial test Seq. | Seq. with low similarity |
|---|---|---|
| 1 | 2412 | 707 |
| 2 | 6536 | 1844 |
| 3 | 4010 | 1263 |
| 4 | 1847 | 433 |
| 5 | 1041 | 275 |
| 6 | 1418 | 325 |
| 7 | 859 | 153 |
| 1.1.1.1 | 4666 | 578 |
| 2.1.1.37 | 3760 | 1240 |
| 3.2.1.4 | 2876 | 760 |
| 4.1.1.39 | 12614 | 100 |
| 5.4.99.5 | 3596 | 462 |
| 6.3.4.15 | 3400 | 962 |
| 7.2.1.1 | 3392 | 200 |

Table S2: Summary of reduced test sets based on low sequence similarity for families with different EC numbers.

#### S2 Model architecture and keywords encoding

The *Finenzyme* model architecture, described in Figure S1, is similar to the CTRL model and learns the probability of the next amino acid in a sequence based on preceding amino acids and control codes, by training on sequences that include control codes. An example sequence containing $n$ tokens is embedded as $n$ vectors in $\mathbb{R}^d$, where each vector is the sum of a learned token embedding and a sinusoidal positional embedding. These vectors are processed through $l$ layers, each containing two blocks: the first block is a multi-head attention with $h$ heads and a causal mask, while the second block is a feed-forward network with ReLU activation projecting the input to an inner dimension $f$. Layer normalization and residual connections are applied before and after each core function within the blocks. The final token scores, computed from the output of the last layer, are used for either a cross-entropy loss function during training or normalized with softmax during generation. *Finenzyme* differs from the original CTRL by accommodating multiple conditioning tags for generating amino acid sequences. *Finenzyme* has a dimension $d = 1280$, an inner dimension $f = 8192$, 36 layers, and 16 heads per layer, with dropout applied after residual connections. Moreover, the token embeddings layer share the same weights of the final output layer.

The pre-trained encoding vocabulary of the model consists of different categories which are encoded in model tokens identifiers numbered from 0 to 129406. The vocabulary is described in Table S3, and it includes 1) UniProt keywords (uni, 2023), providing a comprehensive set of annotations for various protein functions and properties, 2) Taxonomy NCBI tags (Federhen, 2012), that correspond to different taxonomic classifications, ensuring that the model can account for a wide array of biological diversity, 3) Amino acids, covering the IUPAC amino acids encoding (Pettit and Powell, 2006). Additionally, a special PAD token is employed to pad sequences to a uniform length. During fine-tuning, specific adjustments are made to the vocabulary. Fine-tuning assumes the presence of $k$ clusters, replacing the pre-trained codes with indices ranging from 0 to $k - 1$. Each cluster represents a specific EC number, but also it is possible to define sub-clusters to represent subfamilies. Furthermore, a stop token, which signifies the end of a sequence, is encoded as $k$.

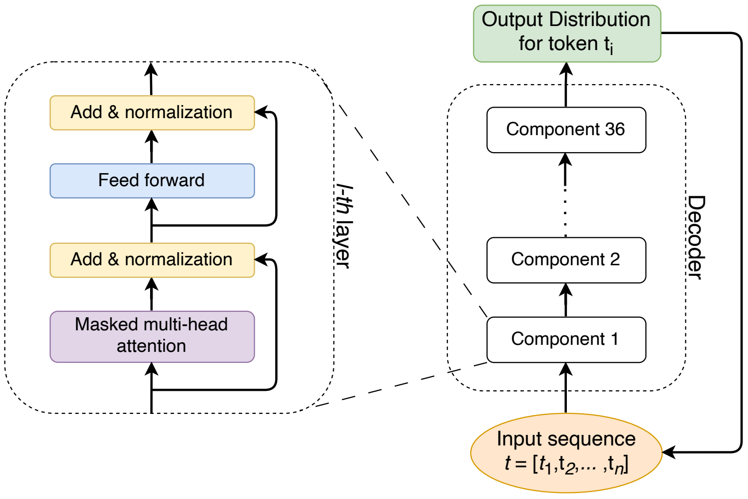


Figure S1: *Finenzyme* model schema, the model has a transformer decoder-only modular architecture.

| Keyword | Encoding |
|---|---|
| UniProt keywords | 0 to 1164 |
| Taxonomy NCBI IDs | 1165 to 129380 |
| Amino acids IUPAC | 129381 to 129405 |
| Pad token | 129406 |

Table S3: Pre-trained model encoding for keywords and amino acids.

#### S3 Test results

The evaluation of the pre-trained and fine-tuned ProGen models was conducted using various metrics, including accuracy, soft accuracy, and perplexity, across different enzyme classes and test setups. Figures S2 and S3 illustrate the performance on general and specific enzyme classes, respectively, with tests categorized by teacher forcing (TF), TF with filtering, prefixed (PF), and PF with filtering. Figures S4–S8 present accuracy, soft accuracy, and perplexity comparisons for high and low-level EC classes under TF testing on both full and filtered test sets. Prefixed testing results for low-level EC classes are shown in Figures S9–S11, comparing performance with a 20 amino acid prefix on full and filtered test sets. Finally, Figures S12 and S13 detail results for the 3.8.1 EC class under both TF and prefixed testing conditions.

**Prefixed (PF) testing can result in high perplexity.** Perplexity results for some sequences generated with PF testing exhibit high values, indicating that these sequences created by the model, whether fine-tuned or not, show low similarity to the natural input sequences, represented by the amino acid prefix. This is not surprising, because PF testing is more challenging, since previous possible errors are not corrected by teacher forcing procedures. Moreover a simple insertion or deletion in the sequence can introduce high perplexity values for all the amino acid predictions downstream of the insertion/deletion itself, even when the downstream prediction are correct instead.

Nonetheless, a portion of the test set demonstrates lower perplexity values. For instance, for the 3.8.1 EC number, 340 out of 1904 tested proteins for the pre-trained model showed 3.86 as mean perplexity, and 1083 out of 1904 tested proteins for the *Finenzyme* model showed an average perplexity of 3.22. Further investigation of sets characterized by high perplexity for the 3.8.1 EC number has been performed with a global alignment for each couple of proteins (true and generated) tested with PF testing, using the Needleman–Wunsch (Needleman and Wunsch, 1970) algorithm. Results (Figure S19) show evidence that *Finenzyme* is generating sequences that, while divergent from the original natural sequences, aligns significantly better to the natural input sequences compared to the pre-trained model generated enzymes (further details in Section S6).

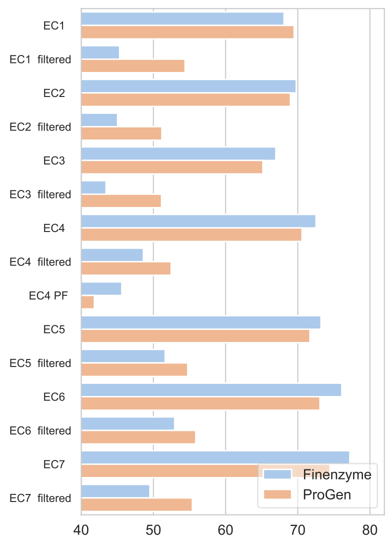


(a)

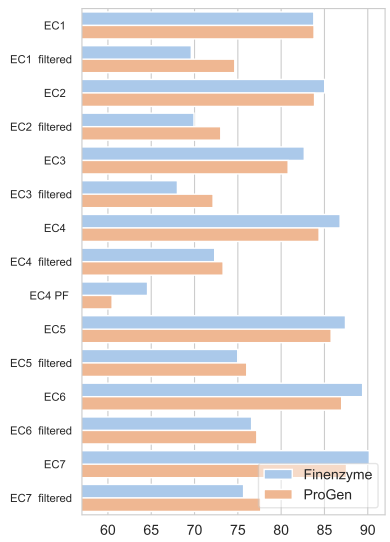


(b)

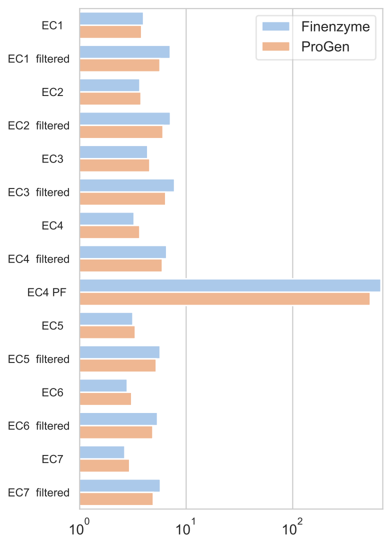


(c)

Figure S2: Performance of Pre-trained and Fine-tuned ProGen Models on general Enzyme Classes Test Data. The graph displays accuracy (a), soft accuracy (b), and perplexity (c) using bar plots for different enzyme classes, which are shown on the x-axis. For each class we group the tests accordingly to the “TF”, “TF filtered”, and “PF”. Blue bars represent fine-tuned results, while orange bars represent pre-trained model results. “TF” stands for teacher forcing, “PF” stands for prefixed, and represents testing without teacher forcing with a prefixed chain of 20 amino acids. Additionally, “filtered” denote whether the test dataset was filtered using BLAST against the training set to retain only sequences with less than 70% identity. Wilcoxon rank sum test shows significance $p < 0.05$ for accuracy and soft accuracy tests without reduced database for five classes, but the p-value raises to $p > 0.7$ for all tests with the reduced database.

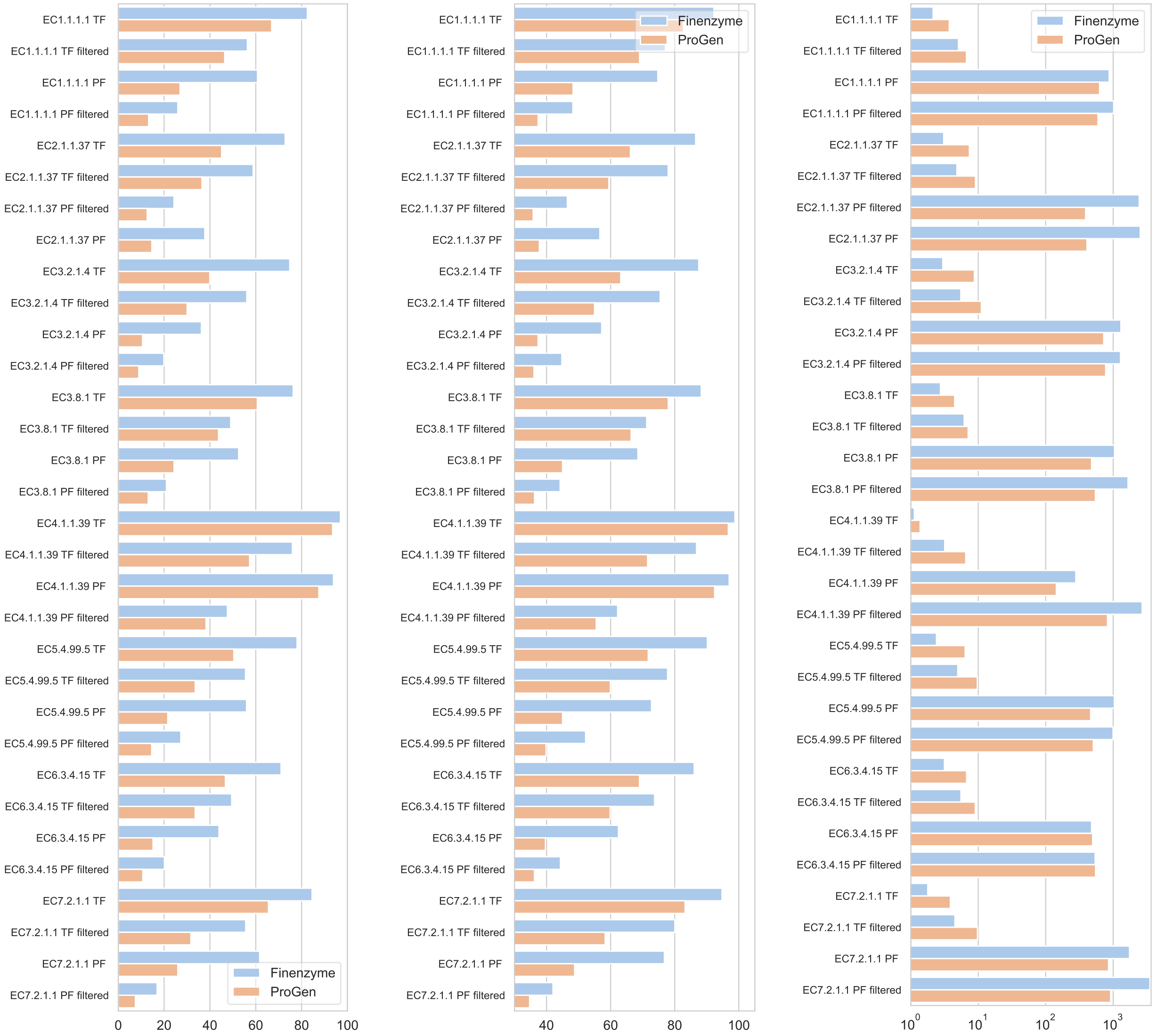


(a) (b) (c)

Figure S3: Performance of Pre-trained and Fine-tuned ProGen Models on specific Enzyme Classes Test Data. The graph displays accuracy (a), soft accuracy (b), and perplexity (c) using bar plots for different enzyme classes, which are shown on the x-axis. For each class we group the tests according to the "TF", "TF filtered", "PF", and "PF filtered". Blue bars represent fine-tuned results, while orange bars represent pre-trained model results. "TF" stands for teacher forcing, "PF" stands for prefixed, and represents testing without teacher forcing with a prefixed chain of 20 amino acids. Additionally, "filtered" denotes whether the test dataset was filtered using BLAST against the training set to retain only sequences with less than 70% identity. Wilcoxon rank sum test has value $p < 10^{-6}$ for PF accuracy and soft accuracy experiments in the barplot, apart from EC4.1.1.39, with $p < 0.04$ for accuracy and $p < 0.07$ for soft accuracy. In the TF setup, for all the metrics, Wilcoxon rank sum test has value $p < 10^{-10}$.

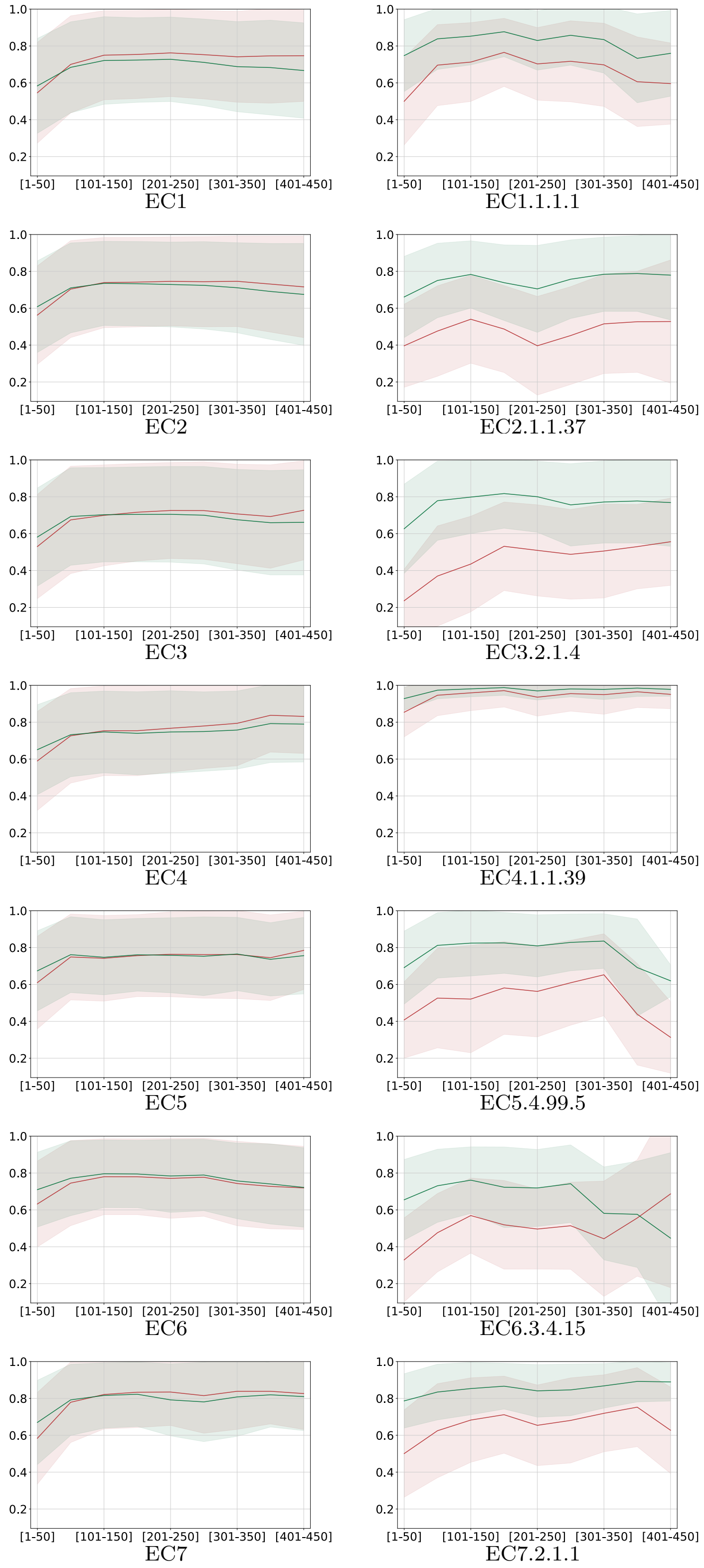


Figure S4: Accuracy comparison of *Finenzyme* and ProGen models on high (left) and low-level (right) EC classes on the full test set. The x-axis reports the position of the amino acid and the y-axis the corresponding average accuracy of the predicted enzyme in the ProGen (red) and *Finenzyme* (green). Shadows represent the standard deviation. The test type is teacher forcing testing.

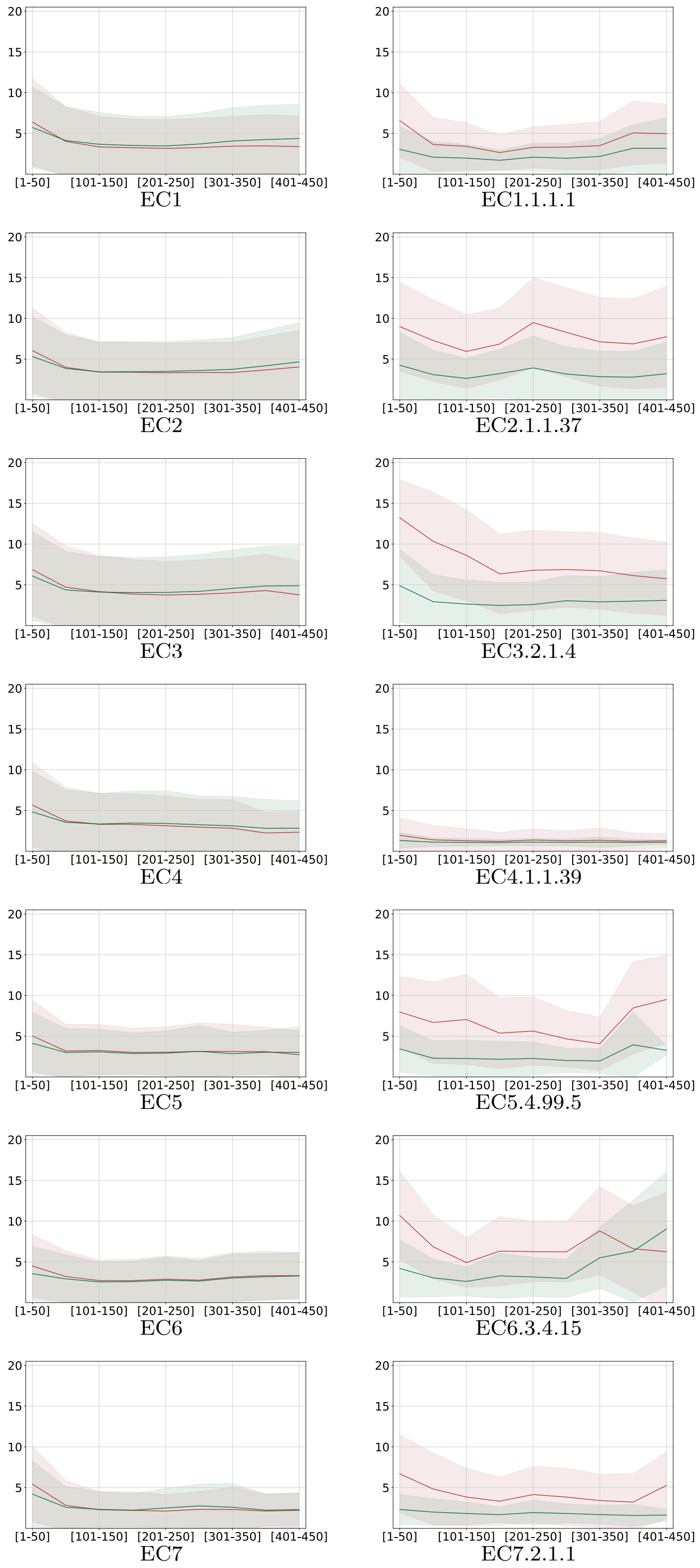


Figure S5: Perplexity comparison of *Finenzyme* and ProGen models on high (left) and low-level (right) EC classes on the full test set. The x-axis reports the position of the amino acid and the y-axis the corresponding average perplexity of the predicted enzyme in the ProGen (red) and *Finenzyme* (green). Shadows represent the standard deviation. The test type is teacher forcing testing.

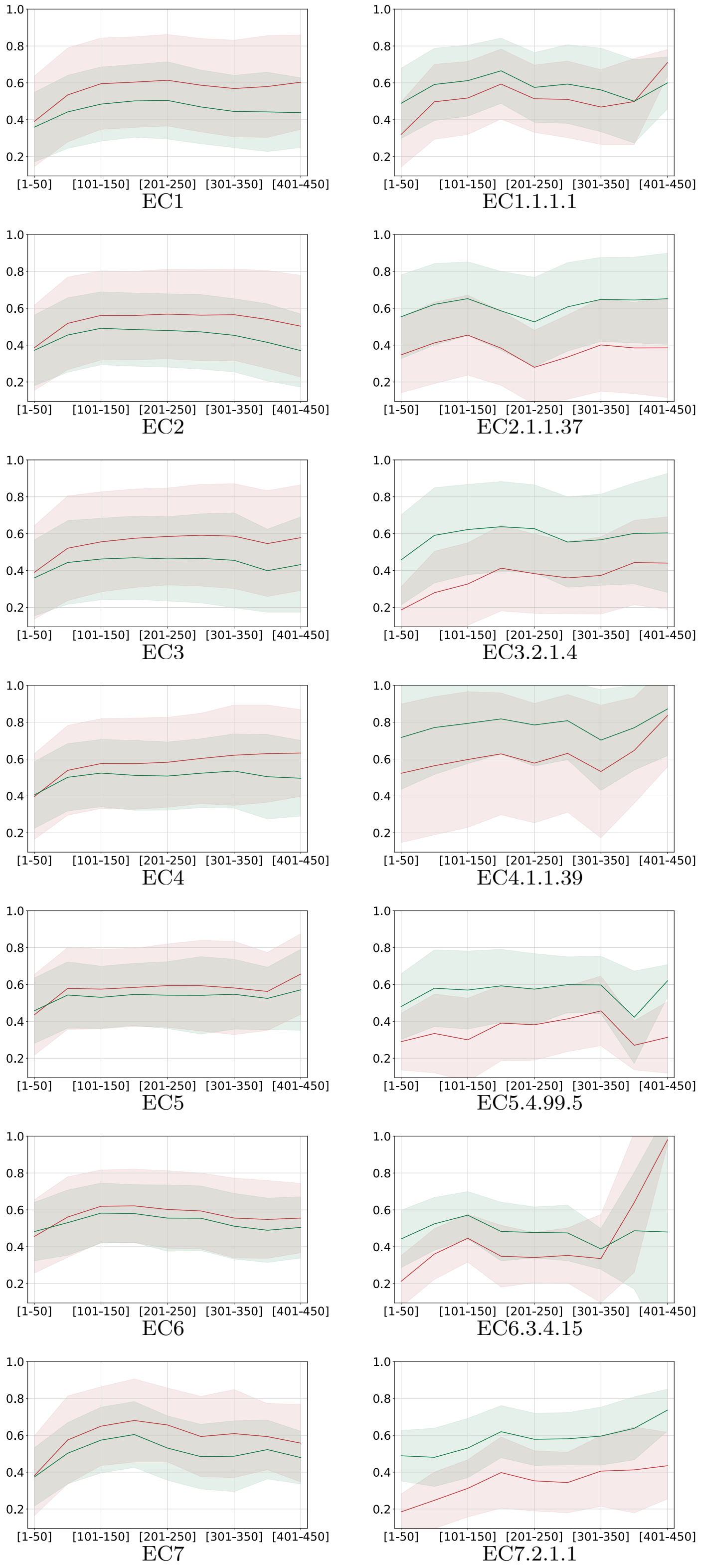


Figure S6: Accuracy comparison of *Finenzyme* and ProGen models on high (left) and low-level (right) EC classes on the filtered test set. The x-axis reports the position of the amino acid and the y-axis the corresponding average accuracy of the predicted enzyme in the ProGen (red) and *Finenzyme* (green). Shadows represent the standard deviation. The test type is teacher forcing testing.

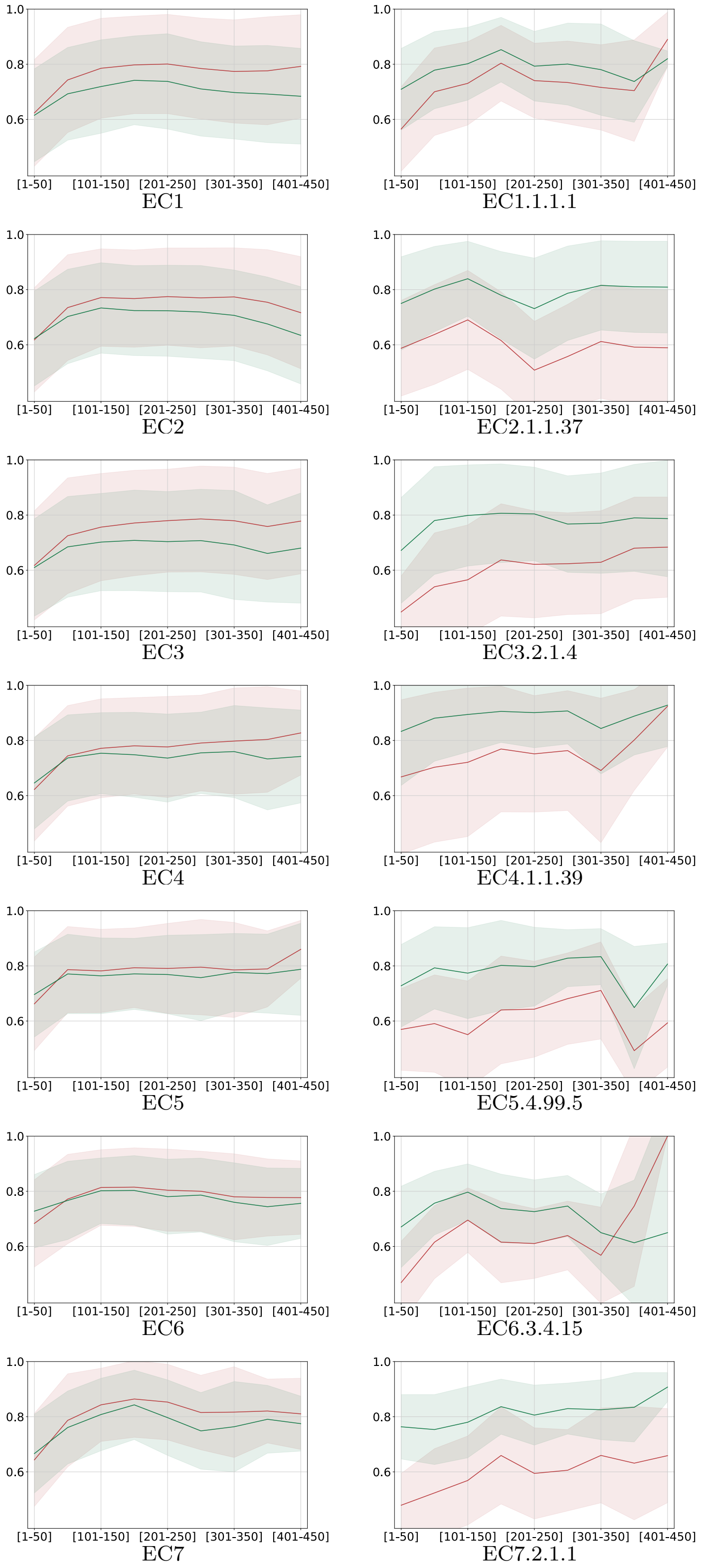


Figure S7: Soft accuracy comparison of *Finenzyme* and ProGen models on high (left) and low-level (right) EC classes on the filtered test set. The x-axis reports the position of the amino acid and the y-axis the corresponding average soft accuracy of the predicted enzyme in the ProGen (red) and *Finenzyme* (green). Shadows represent the standard deviation. The test type is teacher forcing testing.

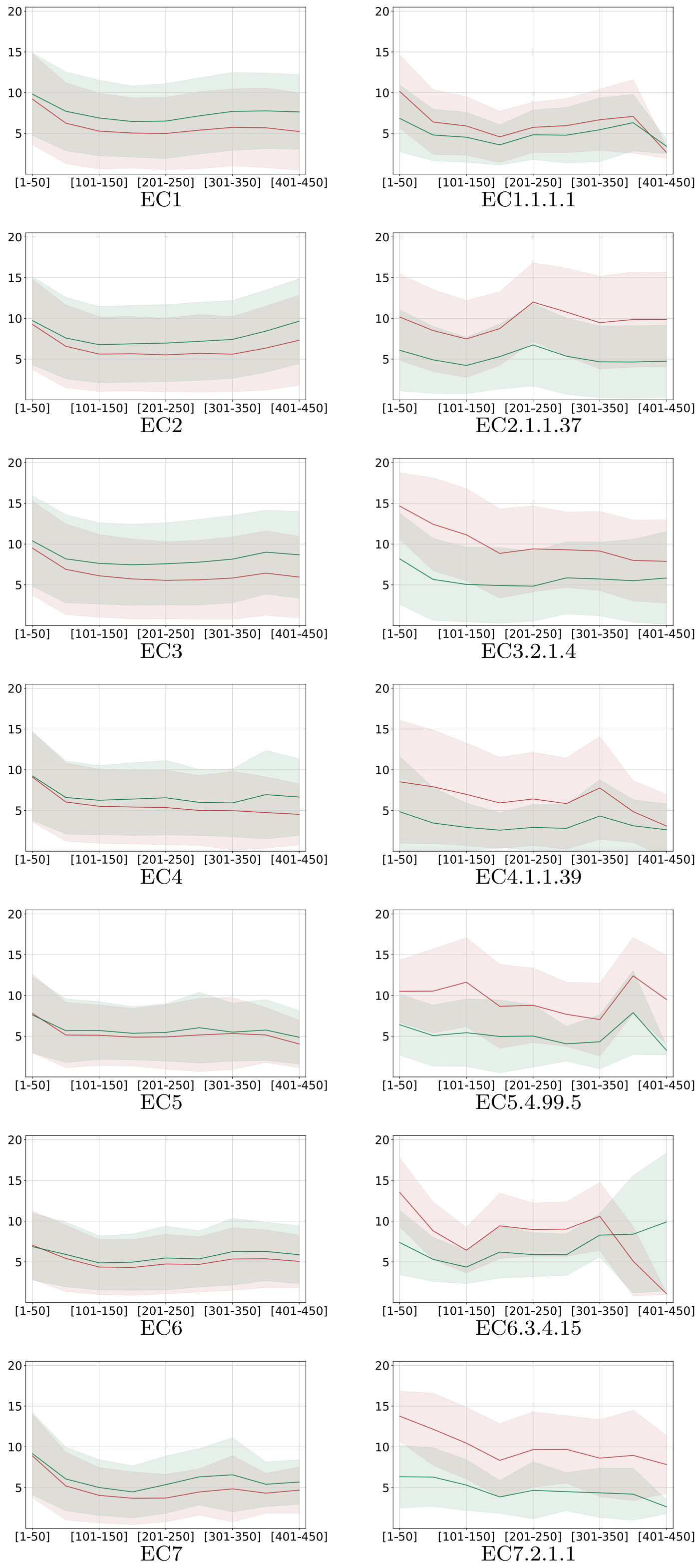


Figure S8: Perplexity comparison of *Finenzyme* and ProGen models on high (left) and low-level (right) EC classes on the filtered test set. The x-axis reports the position of the amino acid and the y-axis the corresponding average perplexity of the predicted enzyme in the ProGen (red) and *Finenzyme* (green). Shadows represent the standard deviation. The test type is teacher forcing testing.

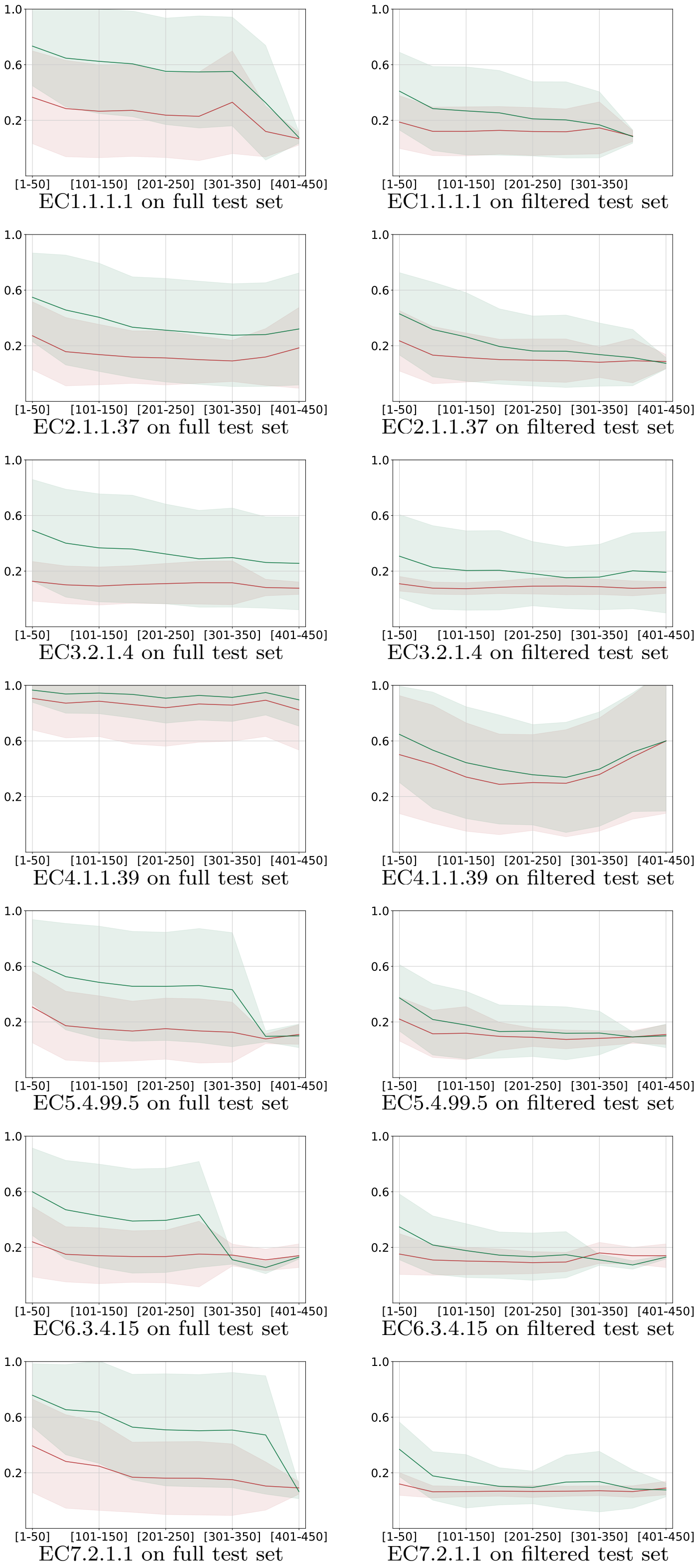


Figure S9: Accuracy comparison of *Finenzyme* and ProGen models on full test (left) and filtered test (right), on low-level EC classes. The x-axis reports the position of the amino acid and the y-axis the corresponding average accuracy of the predicted enzyme in the ProGen (red) and *Finenzyme* (green). Shadows represent the standard deviation. The test type is prefixed testing, with prefix of 20 amino acids.

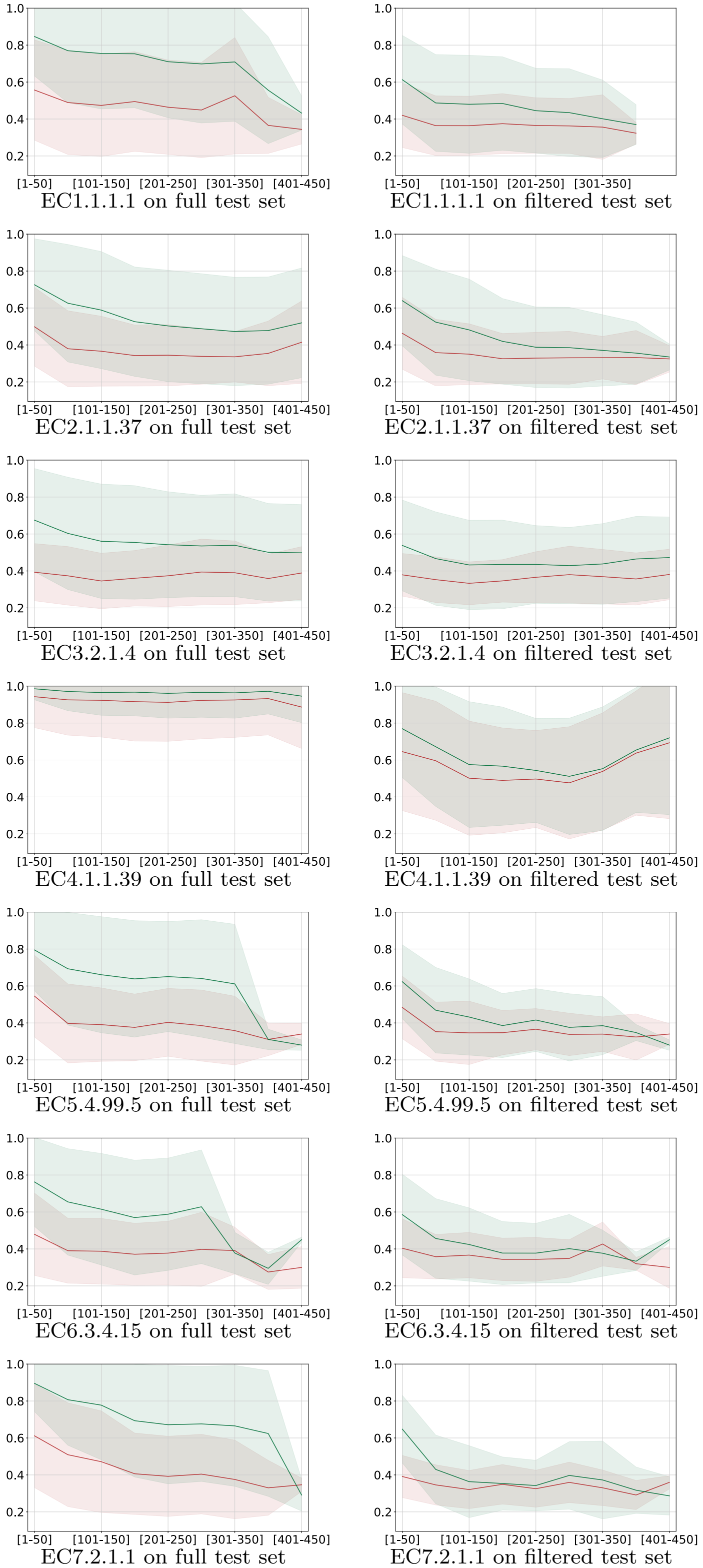

EC1.1.1.1 on full test set
EC1.1.1.1 on filtered test set
EC2.1.1.37 on full test set
EC2.1.1.37 on filtered test set
EC3.2.1.4 on full test set
EC3.2.1.4 on filtered test set
EC4.1.1.39 on full test set
EC4.1.1.39 on filtered test set
EC5.4.99.5 on full test set
EC5.4.99.5 on filtered test set
EC6.3.4.15 on full test set
EC6.3.4.15 on filtered test set
EC7.2.1.1 on full test set
EC7.2.1.1 on filtered test set

Figure S10: Soft accuracy comparison of *Finenzyme* and ProGen models on full test (left) and filtered test (right), on low-level EC classes. The x-axis reports the position of the amino acid and the y-axis the corresponding average soft accuracy of the predicted enzyme in the ProGen (red) and *Finenzyme* (green). Shadows represent the standard deviation. The test type is prefixed testing, with prefix of 20 amino acids.

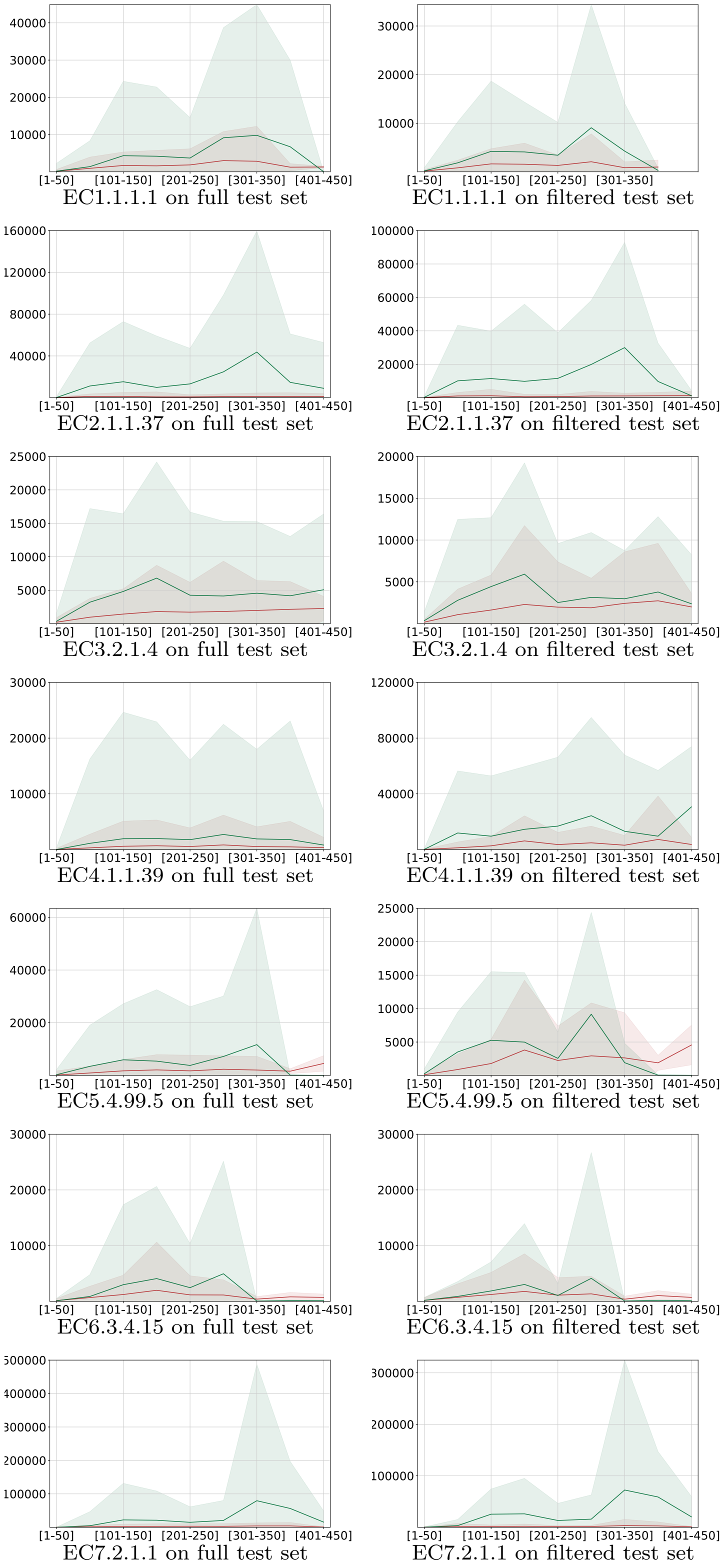


Figure S11: Perplexity comparison of *Finenzyme* and ProGen models on full test (left) and filtered test (right), on low-level EC classes. The x-axis reports the position of the amino acid and the y-axis the corresponding average perplexity of the predicted enzyme in the ProGen (red) and *Finenzyme* (green). Shadows represent the standard deviation. The test type is prefixed testing, with prefix of 20 amino acids.

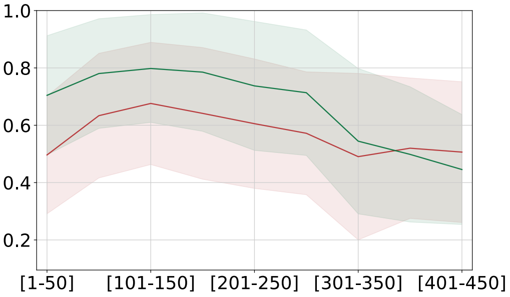


Accuracy on full test set

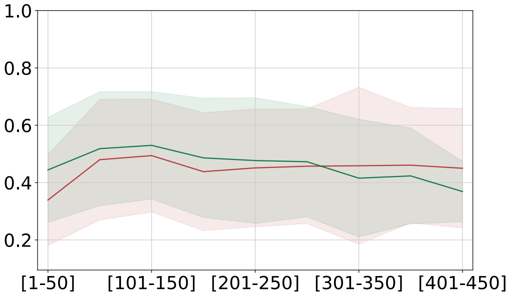


Accuracy on filtered test set

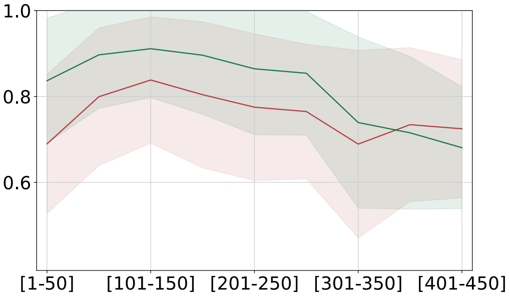


Soft accuracy on full test set

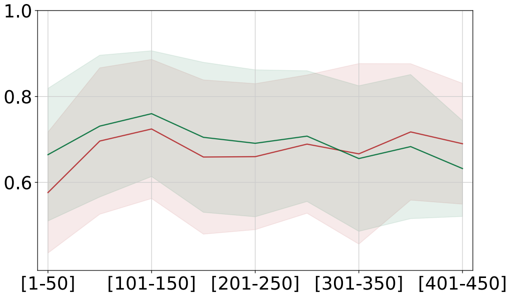


Soft accuracy on filtered test set

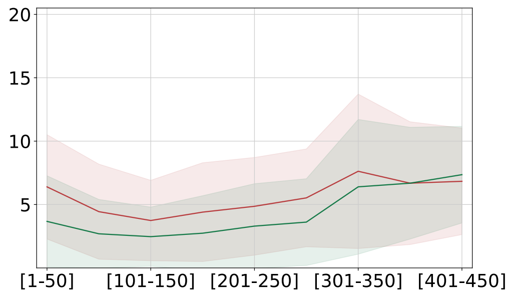


Perplexity on full test set

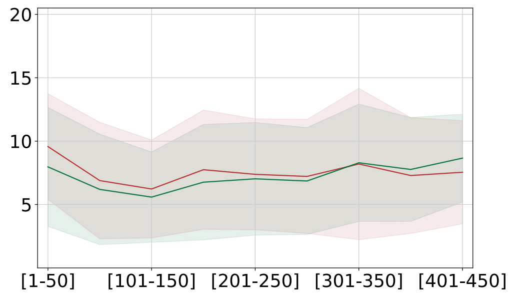


Perplexity on filtered test set

Figure S12: Accuracy, soft accuracy and perplexity comparison of *Finenzyme* and ProGen models on full test (left) and filtered test (right), on the 3.8.1 EC class. The x-axis reports the position of the amino acid and the y-axis the corresponding average metric of the predicted enzyme in the ProGen (red) and *Finenzyme* (green). Shadows represent the standard deviation. The test type is teacher forcing testing.

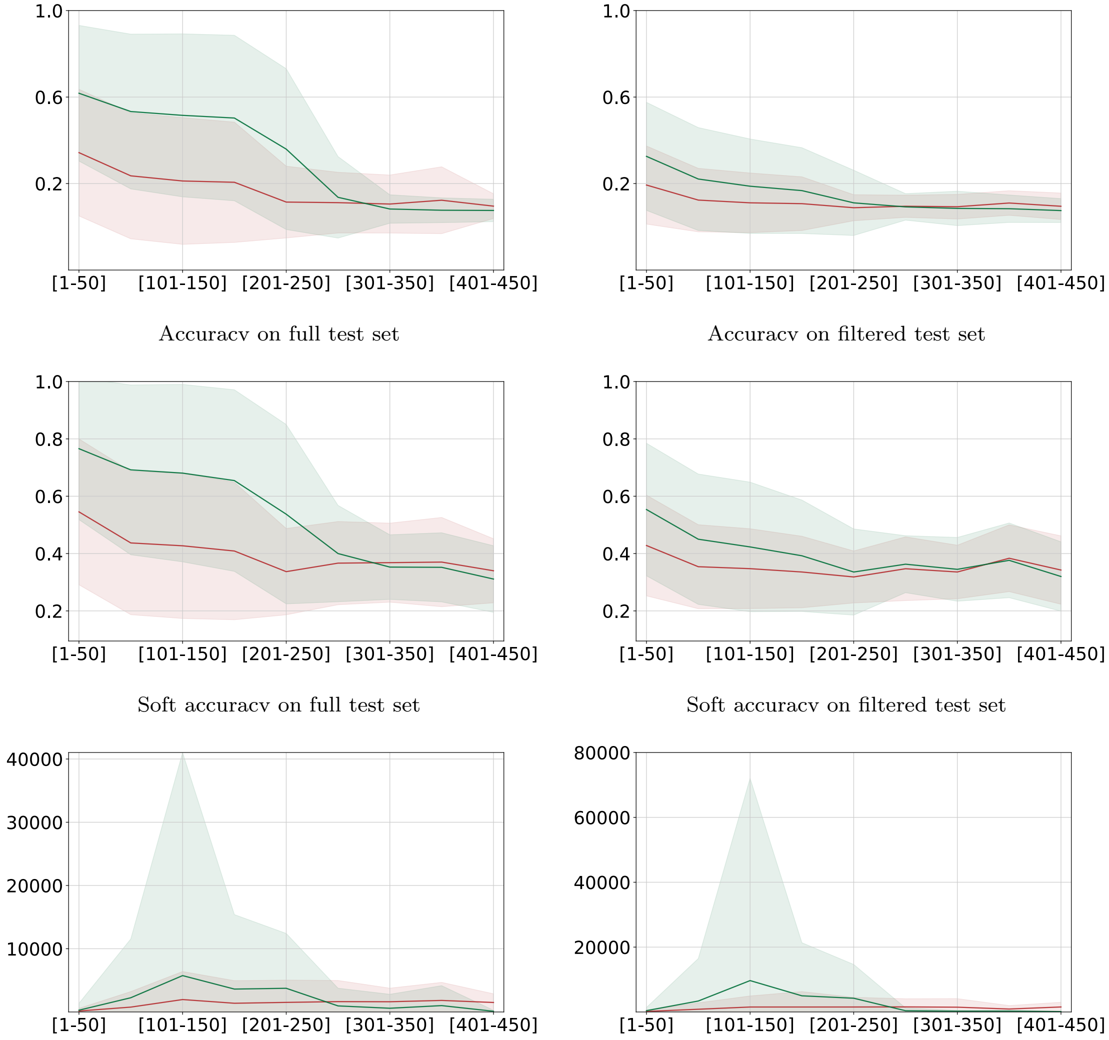


Accuracy on full test set

Accuracy on filtered test set

Soft accuracy on full test set

Soft accuracy on filtered test set

Perplexity on full test set

Perplexity on filtered test set

Figure S13: Accuracy, soft accuracy, and perplexity comparison of *Finenzyme* and ProGen models on full test (left) and filtered test (right), on the 3.8.1 EC class. The x-axis reports the position of the amino acid and the y-axis the corresponding average metric of the predicted enzyme in the ProGen (red) and *Finenzyme* (green). Shadows represent the standard deviation. The test type is prefixed testing, with a prefix of 20 amino acids.

### S4 Only-keywords generation, CLEAN results, and per-class analysis of ESMFold predictions

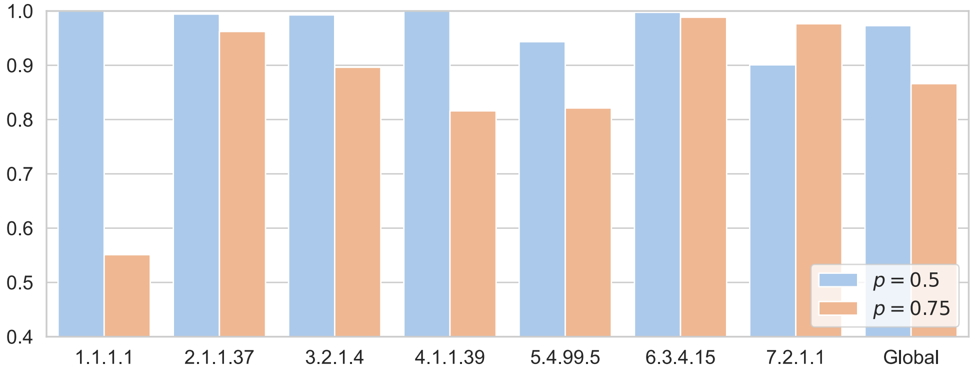


Figure S14: Recall metric for CLEAN EC number predictions. Bar plots show the recall scores for EC number predictions for the generated enzymes, with blue bars representing $p = 0.50$ and red bars representing $p = 0.75$. "Global" represents the weighted average recall score when considering all EC numbers combined.

In this section, we present an analysis of the generated enzymes, examining 1) CLEAN EC number predictions starting from primary structure, and 2) relationships between structural similarity, sequence similarity, and ESMFold prediction confidence for proteins of specific EC classes generated from scratch using keywords only. Sequence and structural similarity are computed through Foldseek.

Table S4 summarizes the distribution of duplicates between natural and *Finenzyme* enzymes generated using different top-$p$ sampling parameters ($p = 0.5$ and $p = 0.75$). Each row represents a specific EC class, showing the number of duplicate sequences generated out of 1000 generated sequences.

Figure S14 presents CLEAN prediction results in terms of recall measures, for each specific EC number considered and for all the EC numbers combined, starting from generated sequences after removing the duplicates described in Table S4.

Figures S15 and S16 show the correlation between structural similarity (TM-score) and sequence identity, as well as the correlation between ESMFold prediction confidence (pLDDT) and structural similarity (TM-score) to the best match found between *Finenzyme* generated and known proteins in the PDB.

| EC | Generated $p = 0.5$ duplicates | Generated $p = 0.75$ duplicates |
|---|---|---|
| 1.1.1.1 | 999 | 285 |
| 2.1.1.37 | 302 | 90 |
| 3.2.1.4 | 863 | 336 |
| 4.1.1.39 | 982 | 641 |
| 5.4.99.5 | 682 | 79 |
| 6.3.4.15 | 610 | 135 |
| 7.2.1.1 | 748 | 363 |

Table S4: Duplicates distribution over the specific EC classes, regarding the enzyme generation by scratch using keywords only. Each generation round with a specific $p$ parameter of top-$p$ sampling counts 1000 generated sequences.

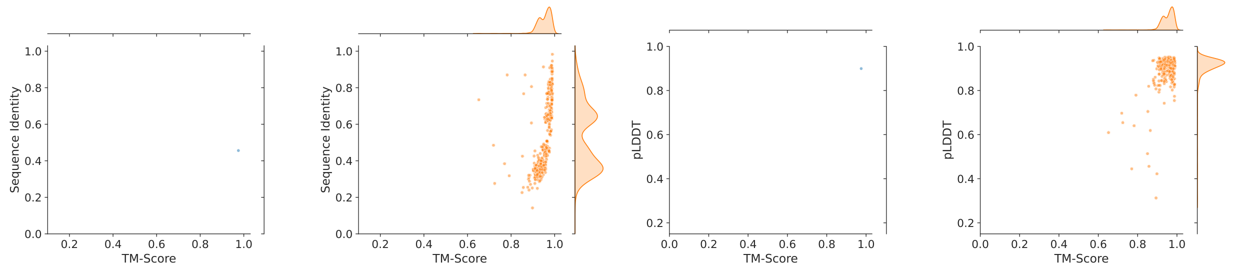


(a) EC1.1.1.1 (b) EC1.1.1.1 (c) EC1.1.1.1 (d) EC1.1.1.1

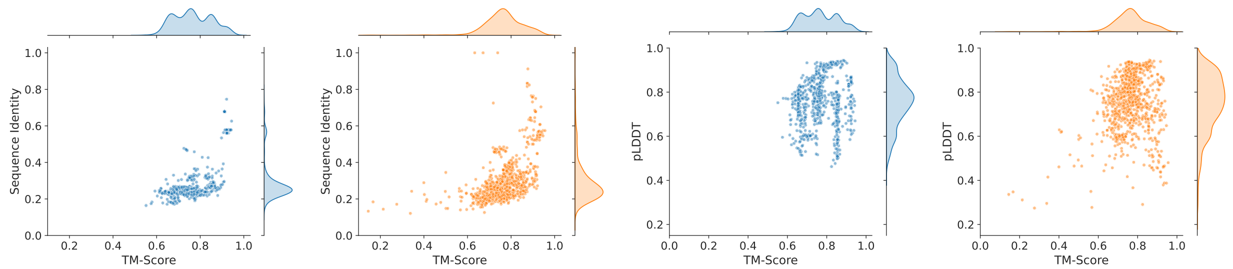


(a) EC2.1.1.37 (b) EC2.1.1.37 (c) EC2.1.1.37 (d) EC2.1.1.37

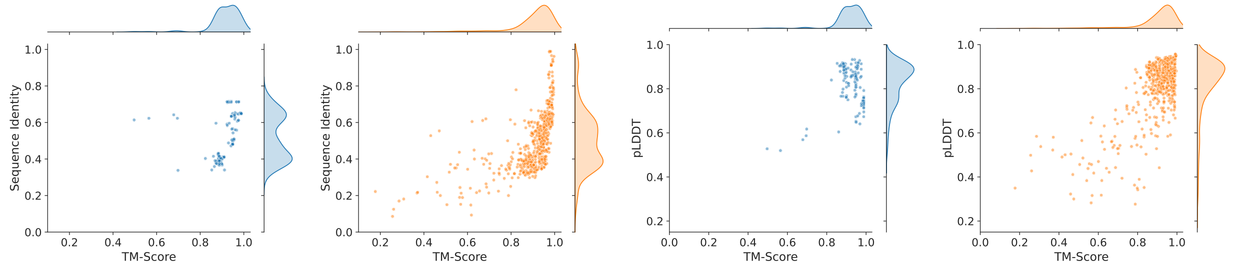


(a) EC3.2.1.4 (b) EC3.2.1.4 (c) EC3.2.1.4 (d) EC3.2.1.4

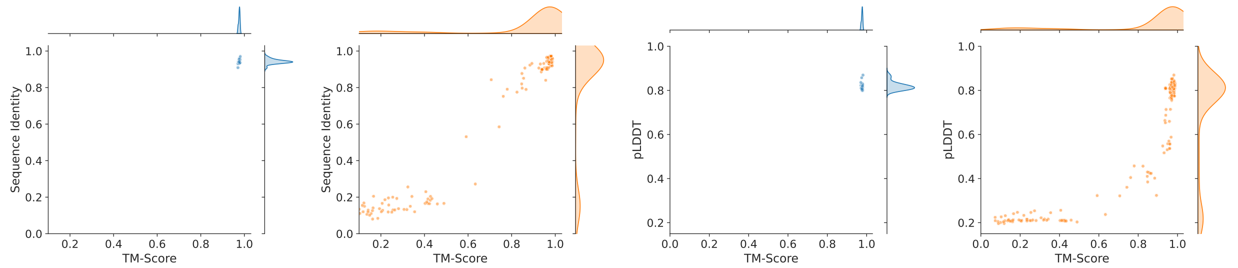


(a) EC4.1.1.39 (b) EC4.1.1.39 (c) EC4.1.1.39 (d) EC4.1.1.39

Figure S15: *Finenzyme* generated enzymes: relationships between structural and sequential similarity with natural enzymes and ESMFold prediction confidence. (a) and (b): Correlation between structural similarity (TMscore) and sequence identity (BLAST Max ID) between natural and generated enzymes across predicted structures from low-level EC classes. (c) and (d): Correlation between ESMFold prediction confidence (pLDDT) and structural similarity to known proteins in the PDB (TMscore). Blue scatterplots refer to top-$p = 0.5$ (a and c), orange to top-$p = 0.75$ (b and d) nucleus filtering. Distributions on top and on the right of each subfigure refers to Sequence identity, TM-score and pLDDT values across the predicted structures from 1.1.1.1, 2.1.1.37, 3.2.1.4, and 4.1.1.39 low-level EC categories.

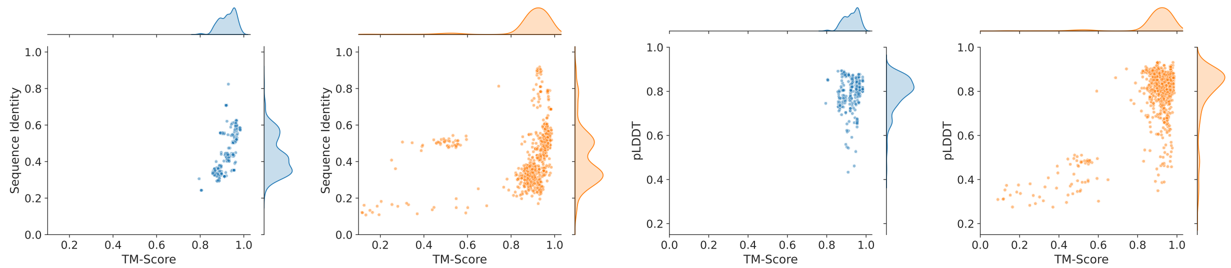


(a) EC5.4.99.5 (b) EC5.4.99.5 (c) EC5.4.99.5 (d) EC5.4.99.5

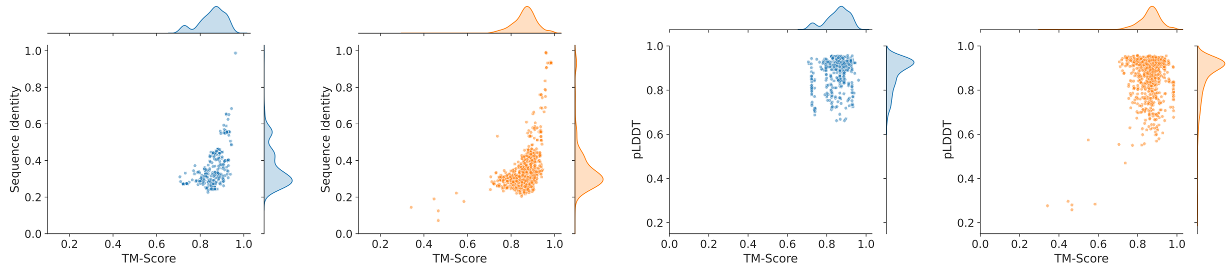


(a) EC6.3.4.15 (b) EC6.3.4.15 (c) EC6.3.4.15 (d) EC6.3.4.15

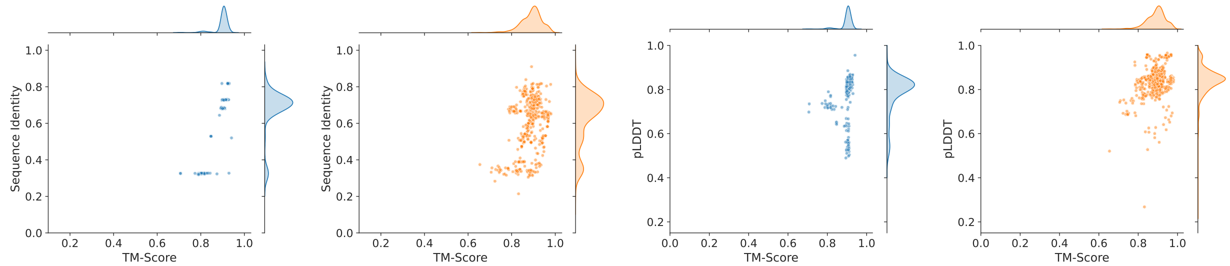


(a) EC7.2.1.1 (b) EC7.2.1.1 (c) EC7.2.1.1 (d) EC7.2.1.1

Figure S16: *Finenzyme* generated enzymes: relationships between structural and sequential similarity with natural enzymes and ESMFold prediction confidence. (a) and (b): Correlation between structural similarity (TMscore) and sequence identity (BLAST Max ID) between natural and generated enzymes across predicted structures from low-level EC classes. (c) and (d): Correlation between ESMFold prediction confidence (pLDDT) and structural similarity to known proteins in the PDB (TMscore). Blue scatterplots refer to top-$p = 0.5$ (a and c), orange to top-$p = 0.75$ (b and d) nucleus filtering. Distributions on top and on the right of each subfigure refers to Sequence identity, TM-score and pLDDT values across the predicted structures from 5.4.99.5, 6.3.4.15, and 7.2.1.1 low-level EC categories.

### S5 Prefixed generation and directed evolution

In this section, we present an analysis of the generated sequences using both keywords and an amino acid prefix using three particular protein targets as input to *Finenzyme*. The objective of this task is to reproduce in-silico the "Directed Evolution" of a protein target, in order to enhance its catalytic activity. Our primary focus is on three sequences of significant scientific interest: alcohol dehydrogenase (EC 1.1.1.1: P11766): Known for its high activity towards formaldehyde. DNA repair enzyme (EC 2.1.1.37: P05102): Prevents the incorporation of damaged bases into DNA, maintaining genomic stability and integrity. Metabolic enzyme (EC 6.3.4.15: Q9SL92): A potential target for genetic manipulation to improve crop yield and stress tolerance through enhanced metabolic efficiency.

The generation process, detailed in the main paper Section 4.5, produced a total of 3000 sequences. First, duplicate sequences were filtered out, and the count of unique sequences per target is provided in Table S5. Next, predicted structures were obtained using ESMfold and evaluated versus the target enzymes and also searched against the Protein Data Bank (PDB) using Foldseek. Structures with a pLDDT greater than 0.7, indicating a good confidence score as defined by ESM authors, were retained. Those with a TM-score to the target greater than 0.7 and alignment coverage greater than 0.7 were selected. These parameters can be selected according to the specific objective of the enzyme generation. For instance, to better guarantee a large structural similarity with the target enzyme, we can enlarge the thresholds for the TM-score and the alignment coverage.

The entire filtering process and data cardinality are detailed in Table S5.

We used the Foldseek representation of enzymes and the MMseqs suite, optimized for the "3D" alphabet of Foldseek, to cluster the filtered sequences (Hauser et al., 2016). From each cluster, a representative enzyme was chosen, and multiple sequence alignments were performed incorporating the natural enzyme targets. These results are displayed in a circular tree visualization. Figs. S17 and S18 present circular trees for the directed evolution of enzymes P11766 and Q9SL92.

Table S6 describes the substrate–enzyme interaction data available in UniProt for the chosen targets, and UniKP prediction on both canonical and isomeric substrate representations. Starting from these values, we predicted $k_m$ for each of these substrate–enzyme couples, as detailed in Section 4.8 of the paper.

| **Model** | **Protein** | **Unique seq.** | **PDB hits** | **Target hit** | **pLDDT>0.7** | **TMscore>0.7 (target)** | **Coverage>0.7 (target)** |
|---|---|---|---|---|---|---|---|
| 1.1.1.1 | P11766 | 402 / 452 | 402 / 359 | 401 / 2 | 392 / 2 | 392 / 2 | 83 / 2 |
| 2.1.1.37 | P05102 | 475 / 479 | 471 / 475 | 471 / 472 | 462 / 274 | 460 / 271 | 136 / 140 |
| 6.3.4.15 | Q9SL92 | 462 / 389 | 443 / 379 | 318 / 357 | 107 / 221 | 106 / 130 | 78 / 76 |

Table S5: Prefixed generation filtering process analysis. Results are presented in the form directed set / inverted resulting set in terms of cardinality of the generated proteins resulting after filtering.

| **ID** | **Predicted ($k_m$) canonical/isomeric** | **Substrate** |
|---|---|---|
| P11766 | 57.143/34.503 $\mu$M | 20-HETE (cid 5283157) |
| P05102 | -/23.945 $\mu$M | DNA (CHEBI:85452) |
| P05102 | 15.302/18.683 $\mu$M | S-adenosyl-L-methionine (cid 24762165) |
| Q9SL92 | 72.352/64.913 $\mu$M | biotin (cid 6560210) |
| Q9SL92 | 63.249/66.443 $\mu$M | ATP (cid 5461108) |
| Q9SL92 | 52.173/- $\mu$M | methylcrotonoyl-CoA carboxylase (cid 29969) |

Table S6: Substrate–enzyme interaction data from UnipProt. The predicted $k_m$ kinetic values are UniKP predictions of the interaction between the natural target and the substrate. SMILEs have been retrieved from `https://pubchem.ncbi`, apart from the DNA SMILE, that have been retrieved from `https://www.ebi.ac.uk/chebi`.

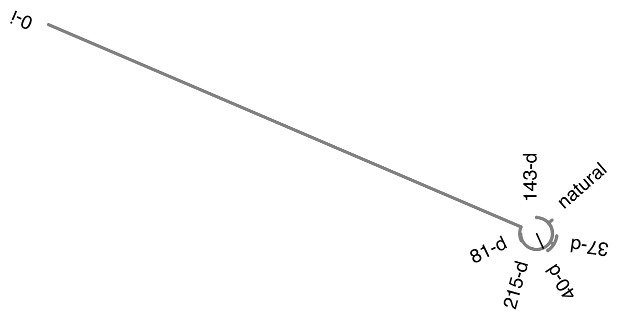


Figure S17: *Finenzyme* directed evolution. Circular tree representing the directed evolution of enzyme P11766.

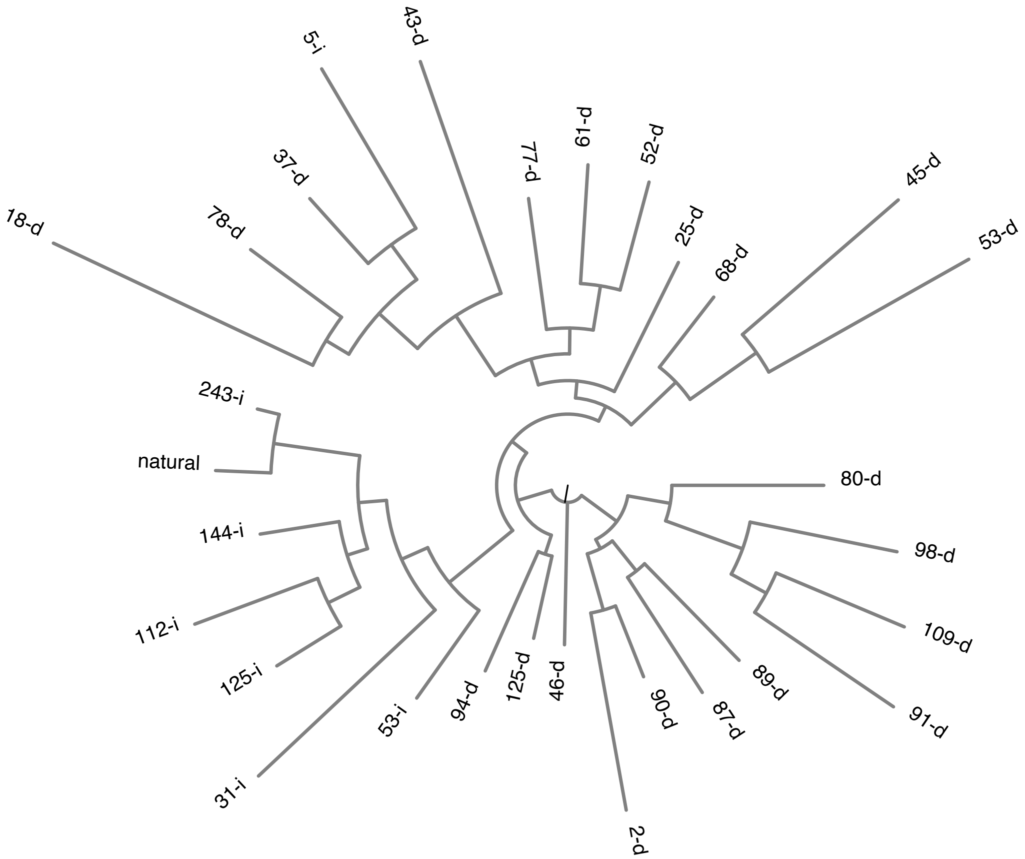


Figure S18: *Finenzyme* directed evolution. Circular tree representing the directed evolution of enzyme Q9SL92.

### S6 *Finenzyme* generation of dehalogenases (EC 3.8.1)

In this section, we provide further information on fine-tuning and sequence generation analysis regarding the family of dehalogenase enzymes. We showed in the main text that the teacher forcing analysis improved model prediction of the natural sequences belonging to the test set. In this case, we also tested whether the model was able to predict better sequences belonging to the test set given a prefix of 20 amino acids only. In this case, we decided to compare ProGen and *Finenzyme* predictions by performing a global sequence alignment through the Needlman-Wunsch algorithm (Needleman and Wunsch, 1970). The latter allows the model to include gaps in case they are needed.

Hyperparameter tuning on gap penalty and gap opening penalty led to the decision of assigning the value -11 to the former and -1 to the latter. Furthermore, the final score was normalized as:

$$\text{Normalized Score} = \frac{\text{Score} - \mu}{\sigma}. \quad (1)$$

The mean and standard deviation were estimated through a Monte Carlo simulation of 100 iterations, where for each alignment the real sequence was shuffled randomly. The differences in performance of ProGen against *Finenzyme* are depicted in Figure S19. A Wilcoxon Signed-Rank Test was performed showing the statistically significant improvement in performance. As we found that *Finenzyme* was a better predictor, we moved onto artificial sequence generation.

In Figure S20 we show the distribution of subclasses distribution. Considering that the cardinality of certain classes was below 1000 sequences, we did not include their keywords, which means that we included keywords only for three subclasses, namely 3.8.1.2, 3.8.1.5 and 3.8.1.3.
After fine-tuning, we generated sequences by providing only the keyword tokens. For instance, when generating sequences belonging to subclass 3.8.1.2, the input to the model was:
`model( <3.8.1.->, <3.8.1.2>, <PAD>, ... ,<PAD>).`
We also tried to generate sequences with only the most general keyword `<3.8.1.->` as input. The results for the generation of enzymes are summarized in Table S7. As nucleus sampling needs some hyperparameter tuning, we found that with top $p = 0.50$ many duplicates where generated for subclass 3.8.1.2, so we also generated enzymes by increasing top $p = 0.75$. As the latter was the most abundant subclass, we realized that the model's entropy for such subclass was lower that for the others (the model was more confident).

As the length of the generated sequences depends on the most probable stop token along the sequence, we tested such heuristic comparing the distribution of the lengths of the generated sequences against the training set. The results of such analysis are depicted in Figure S21. In the figure, the rows show sequences belonging to either only 3.8.1.- (first row) or also to the aforementioned subclasses. The first and second columns are the lengths of the sequences generated with top $p = 0.50$ and top $p = 0.75$ respectively, whereas the third column represents the natural sequences.

We measured maximum sequenced identities through BLAST between generated and natural sequences (Figure S22): we can see that there is a good identity with the training set's sequences, and more constrained sampling ($p = 0.50$) leads to the generation of sequences that, in the majority of the cases, resemble more the training set.

As sequence identity is not as important as structural similarity, we predicted the structure of the generated sequences through ESMFold (Lin et al., 2023). Subsequently, we checked whether the generated structures resemble any experimentally validated structure through Foldseek (Van Kempen et al., 2024). The results for each subclass are depicted in Figure S23, and the general results are depicted in Figure S24. For both $p = 0.50$ and $p = 0.75$ we can see that in general structural similarity is maintained, even when sequence identity drops. Nonetheless, when $p = 0.50$, the average TM-score is 0.92 and the average sequence identity is 0.46. When $p = 0.75$, the average TM-score is 0.68 and the average sequence identity is 0.32. As the TM-score is largely higher than 0.5 in both cases, we can infer that even if some sequences have a low maximum identity, they are still able to fold correctly (Xu and Zhang, 2010).

Fig. S24 compares TM-score results with either the pLDDT or the sequence identity of the highest ranked Foldseek hit per sequence. The pLDDT score is used as a measure of confidence of the predicted structure by ESMFold, and it has been demonstrated that low pLDDT values correspond to wrongly predicted structures (Lin et al., 2023; Jumper et al., 2021).

The Spearman correlation coefficient between pLDDT and TM-score is low when $p = 0.50$ (0.17), but the same coefficient is much higher with $p = 0.75$ ($\rho = 0.76$ and p-value $< 10^{-10}$). These results can justify some of the low TM-scores, that can be due to a wrongly predicted structure by ESMFold rather than a non-meaningful generated sequence.

Interestingly, we noticed that the average sequence identity was always much lower than the average TM-score. When $p = 0.50$, the average TM-score is 0.92 and the average sequence identity is 0.46. When $p = 0.75$, the average TM-score is 0.68 and the average sequence identity is 0.32. The distribution of the TM-score on top of each subfigure (Fig S24) shows that the TM-score is largely higher than 0.5 and this value represents a valuable threshold for considering a pair of proteins in the same fold (Xu and Zhang, 2010). This demonstrates that despite low sequence similarity, the structures are still maintained. Results for each experiment are reported in Figure S22.

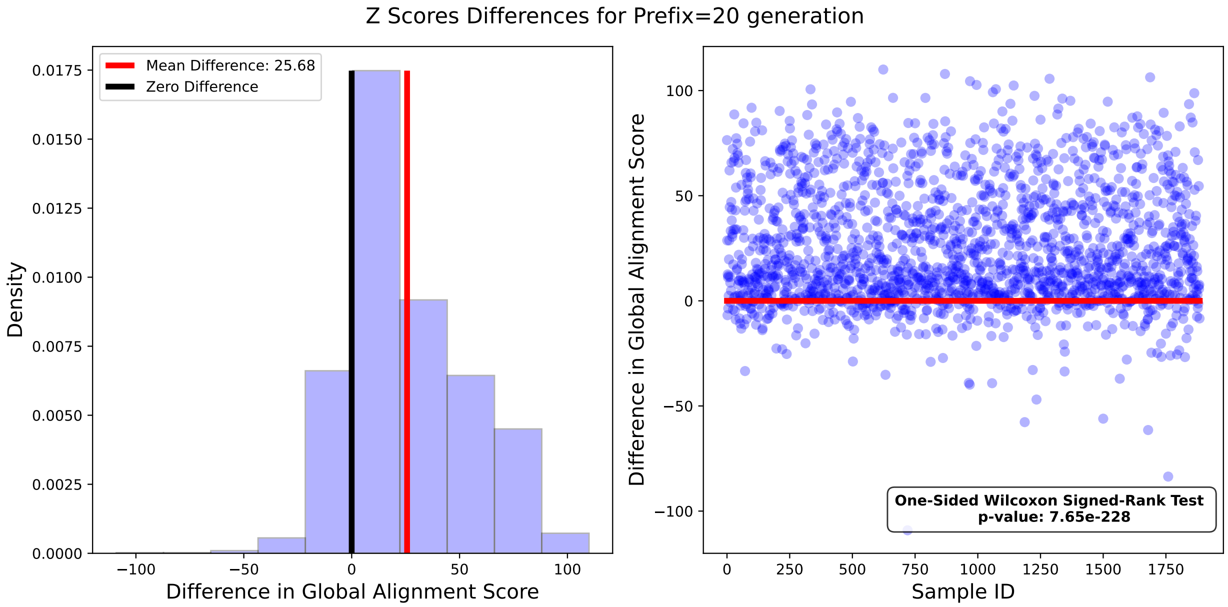


Figure S19: Dehalogenase analysis. On the left, distribution of the differences between *Finenzyme* and ProGen Needleman-Wunsch normalized alignment scores. The Zero black line shows the point where there is no difference in scores. On the right, the red line divides between scores better for *Finenzyme* (above) or better for ProGen (below).

| top-p | EC | Unique Seq. | Dataset Sequences |
|---|---|---|---|
| 0.50 | 3.8.1.- | 611 | 19408 |
| 0.75 | 3.8.1.- | 979 | 19408 |
| 0.50 | 3.8.1.2 | 2 | 11156 |
| 0.75 | 3.8.1.2 | 285 | 11156 |
| 0.50 | 3.8.1.5 | 539 | 5739 |
| 0.75 | 3.8.1.5 | 913 | 5739 |
| 0.50 | 3.8.1.3 | 673 | 1327 |
| 0.75 | 3.8.1.3 | 938 | 1327 |

Table S7: Dehalogenase analysis. Distribution of the number of artificial (third column) and natural (fourth column) dehalogenase sequences belonging to different subclasses.

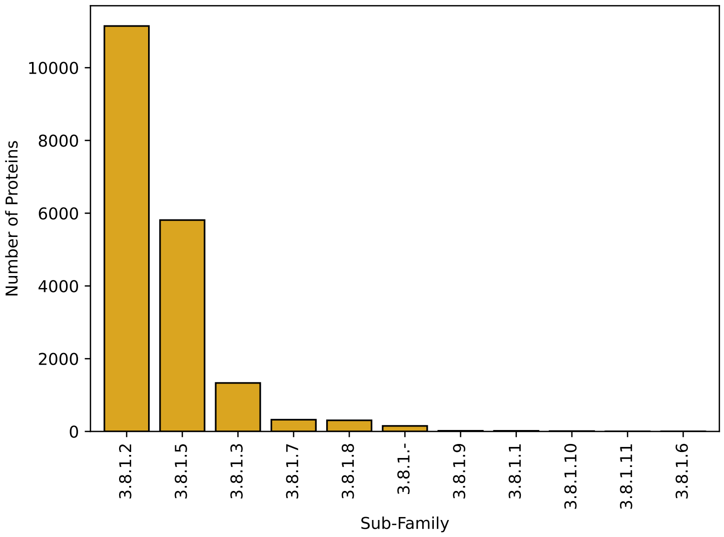


Figure S20: Dehalogenase analysis. Distribution of the number of hydrolases acting on C-halide compounds found in Uniprot.

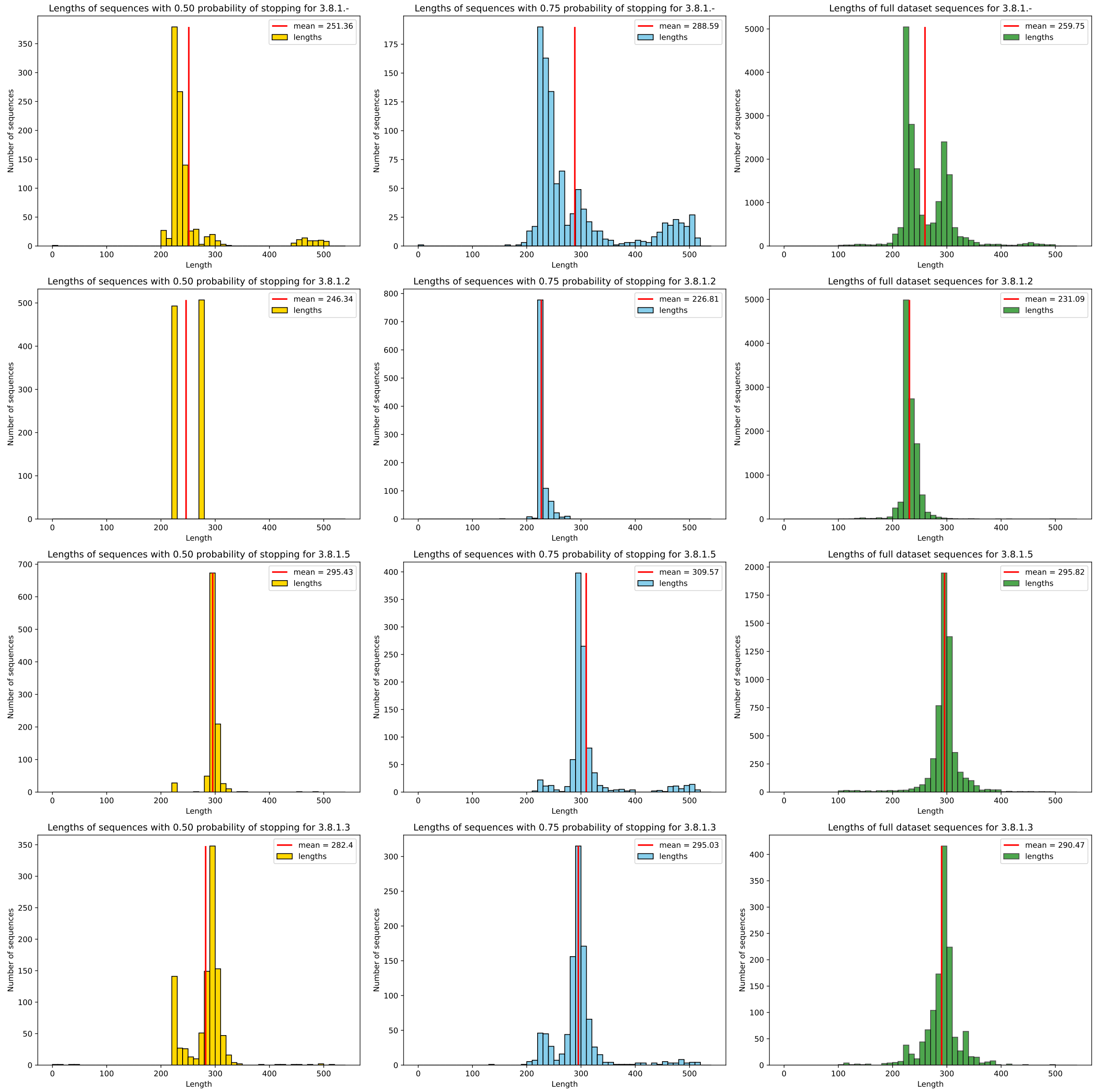


Figure S21: Dehalogenase analysis. Distribution of generated sequences' lengths. The first and second column correspond to the experimental settings, with respectively top-$p$ = 0.50, and top-$p$ = 0.75. The third column represents the distribution of length of the natural enzymes. The first row shows the experiment with no subkeyword prepended, whereas the other 3 rows correspond to experiments with respectively 3.8.1.2, 3.8.1.5 and 3.8.1.3 keywords added.

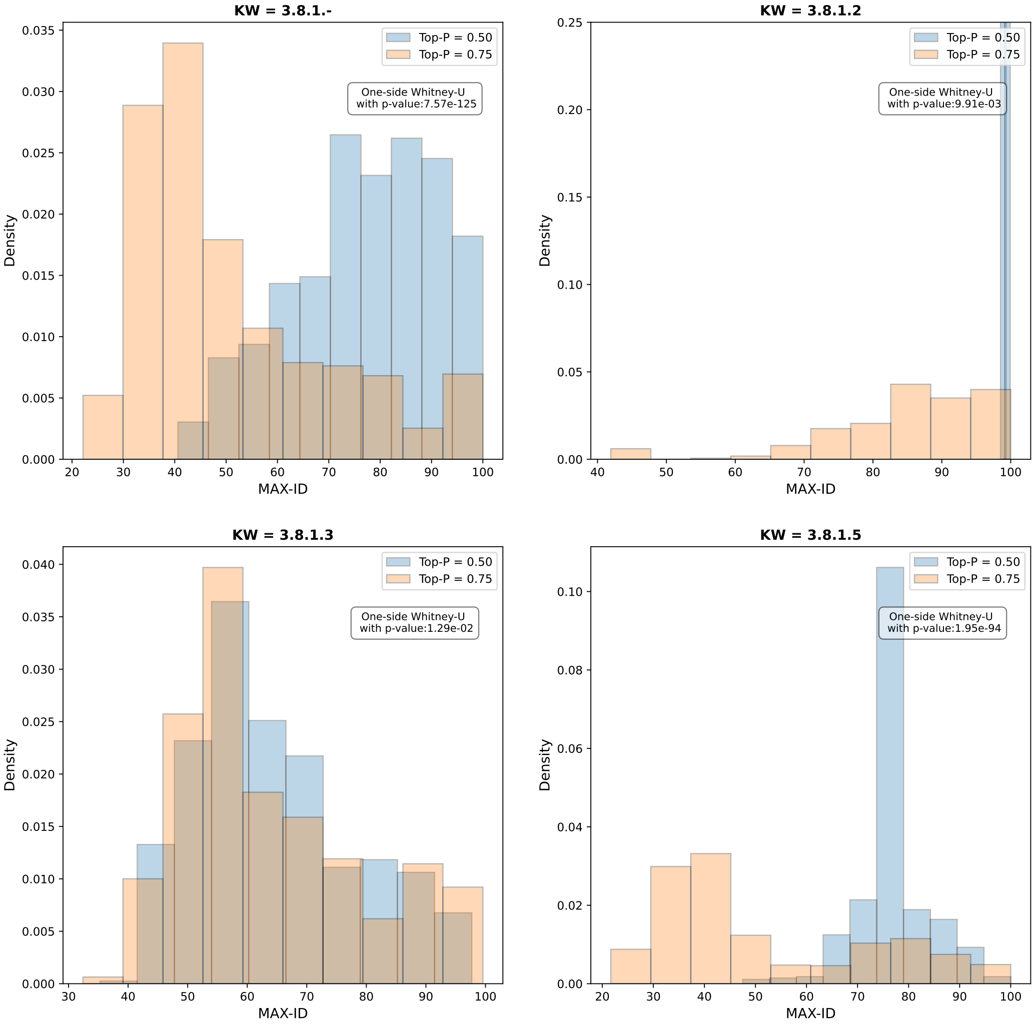


Figure S22: Dehalogenase analysis. Distribution of BLAST MAX-ID hits obtained between generated and natural sequences for EC categories 3.8.1, 3.8.1.2, 3.8.1.3, and 3.8.1.5. Blue bars: top-$p = 0.50$; orange bars: top-$p = 0.75$.

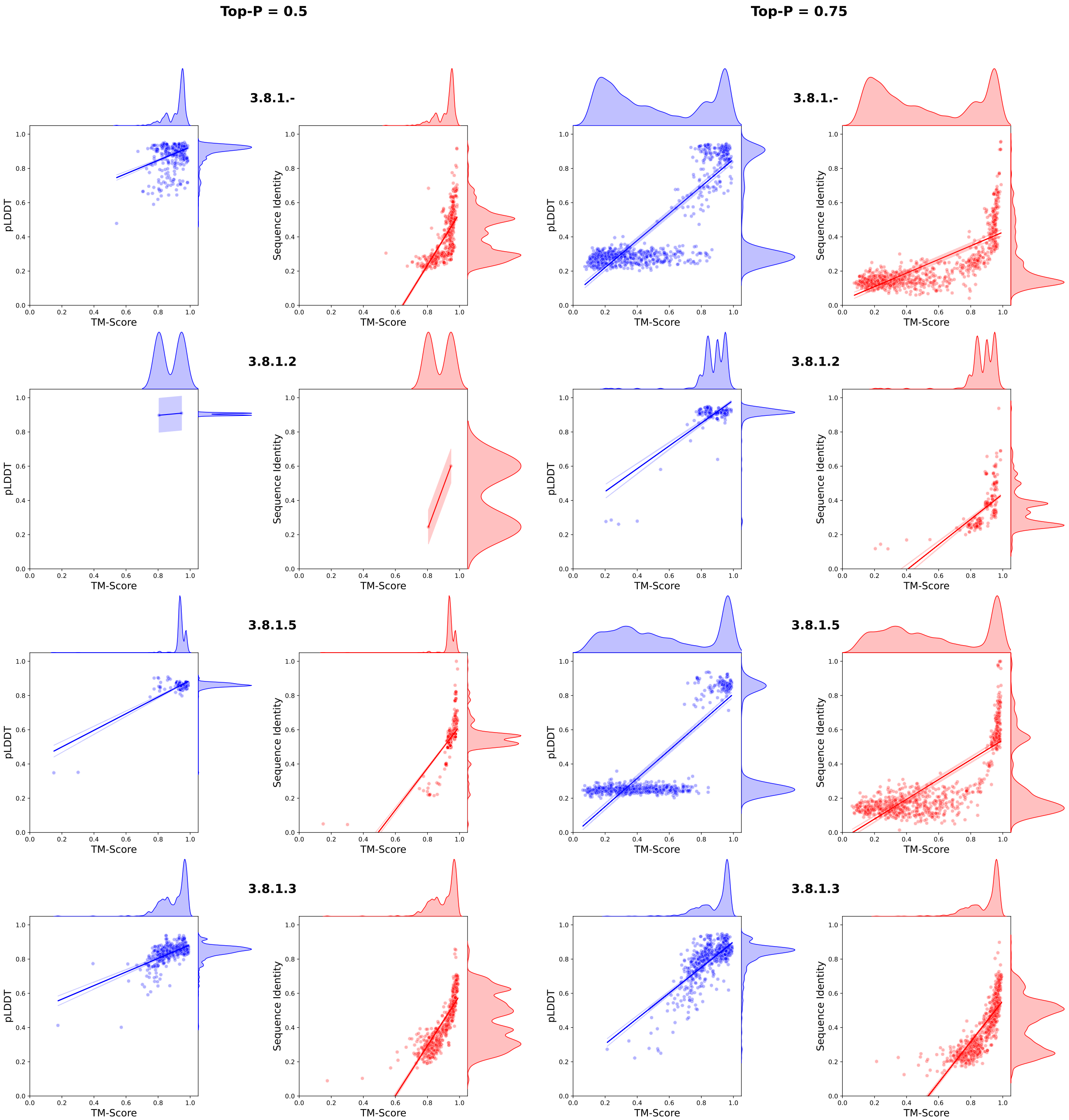


Figure S23: Dehalogenase analysis. Results of Foldseek analysis of the predicted structure for all the generated sequences, distinguished both by the keyword used for the generation and the top-$p$ utilized. Each plot compares the TM-score with either the pLDDT or the Sequence Identity. The straight lines on the plots represent linear fits to display the positive correlations.

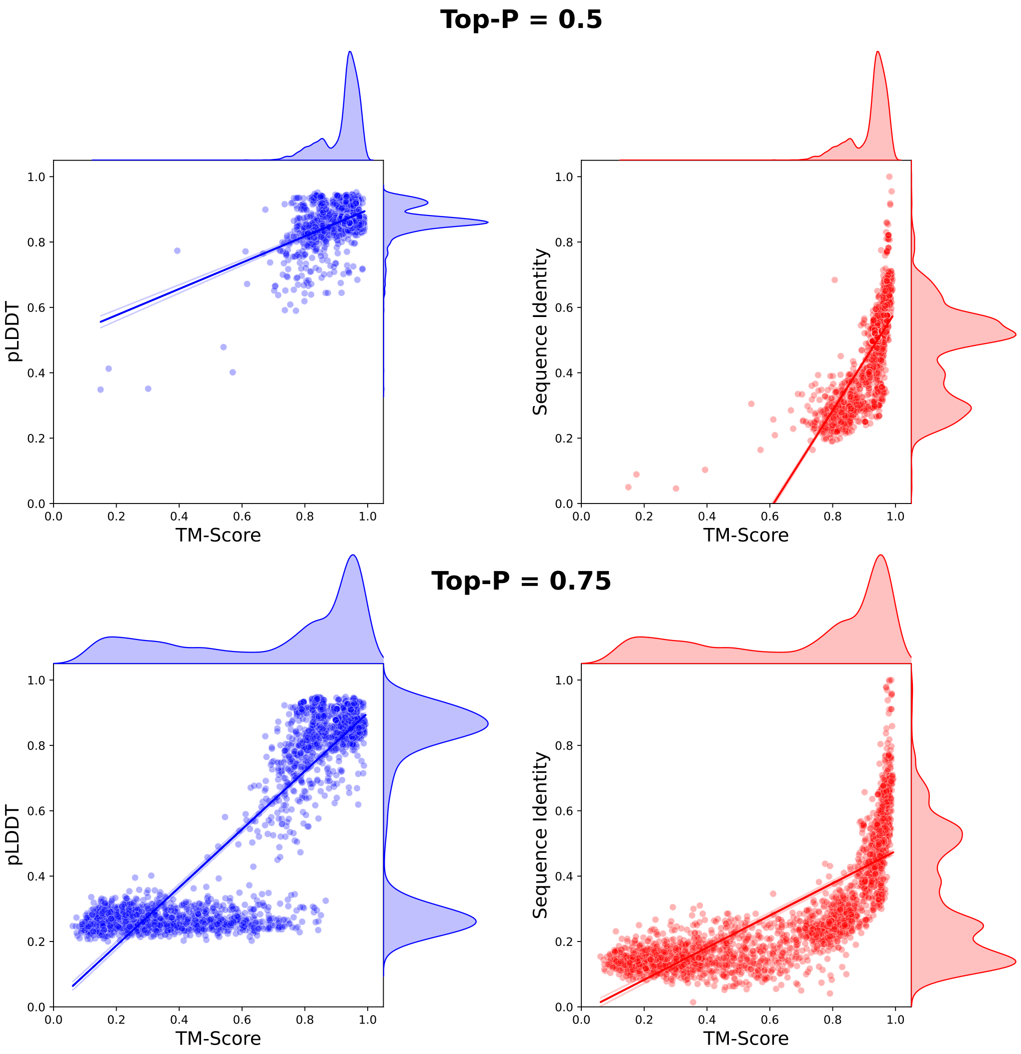


Figure S24: Dehalogenase analysis. Relationships between the TM-score between generated dehalogenases and natural enzymes and pLDDT (left subfigures) and sequence identity (right subfigures). Foldseek has been applied to compute the TM-score. First row: generation with top-$p = 0.5$; second row: generation with top-$p = 0.75$. The straight lines represent linear fits to display the positive correlations. On top and on the right of each subfigure, the distribution of pLDDT, TM-score and sequence identity is displayed.
